# Population genetics of transposable element load: a mechanistic account of observed overdispersion

**DOI:** 10.1101/2020.11.16.382945

**Authors:** Ronald D. Smith, Joshua R. Puzey, Gregory D. Conradi Smith

**Author notes:** These authors contributed equally (Joshua R. Puzey, Gregory D. Conradi Smith).

## Abstract

In an empirical analysis of transposable element (TE) abundance within natural populations of *Mimulus guttatus* and *Drosophila melanogaster*, we found a surprisingly high variance of TE count (e.g., variance-to-mean ratio on the order of 10 to 300). To obtain insight regarding those evolutionary genetic mechanisms that are may underlie the overdispersed population distributions of TE abundance, we developed a mathematical model of TE population genetics that includes the dynamics of element proliferation and purifying selection on TE load. The modeling approach begins with a master equation for a birth-death process and extends the predictions of the classical theory of TE dynamics in several ways. In particular, moment-based analyses of time-dependent and equilibrium population distributions of TE load reveal that overdispersion is likely to arise via copy-and-paste proliferation dynamics, especially when the elementary processes of proliferation and excision are first-order and approximately balanced. Parameter studies and analytic work confirm this result and further suggest that overdispersed population distributions of TE abundance are unlikely to be a consequence of purifying selection on total element load.

## Introduction

The genomics revolution has revealed that a significant portion of eukaryotic genomes consists of transposable elements (TEs, also called mobile DNA elements or transposons). Notable examples include the human and maize genomes, 44% and 85% of which are TE sequences (Mills et al., 2007; Springer et al., 2009). Various mobility mechanisms enable TEs to proliferate and/or change position within a genome. The effect of TEs can range from having little to no consequence on phenotype to being powerful mutagens (Bourque et al., 2018). In addition to the innate tendency of TEs to proliferate, factors such as recombination, epigenetics, and selection contribute to the complex genomic distribution and demography of these elements (Kent et al., 2017). While it is clear that TEs have been an integral part of the long-term evolution of genome architecture, much about the role of TEs in evolution remains unknown. Knowledge of the dynamics of TE abundance in natural populations is an important step toward an increasing understanding of how genomes evolve.

The pioneering and influential population genetic theory of TEs developed by Charlesworth and Charlesworth (1983) used a combination of mathematical analysis, computer simulation, and a limited amount of experimental data, to give theoretical insight into TE dynamics and demographics. This modeling considered a single family of TEs with a drift-diffusion representation of TE proliferation, with either no selection or weak selection acting on total TE copy number. This modeling approach has informed our understanding of the population genetics of TEs for several decades. However, the classical theory does not reproduce experimentally observed within-population variances that often greatly exceed the population mean. The cause of this discrepancy is that the classical modeling approach assumes a binomial distribution of within-population TE loads, which constrains the population variance to be no greater than the population mean.

This paper begins with a brief review of classical TE population genetics. This is followed by an analysis of TE demography derived from genome-sequence data of two natural populations (*Mimulus guttatus* and *Drosophila melanogaster*). Notably, in both cases, we observe that the within-population variance of TE load is highly overdispersed. Because these empirical results violate the predictions of classical TE modeling, we developed a more realistic master equation formulation of the population distribution of TE loads in a large randomly mating population. This alternative population genetic framework simultaneously and self-consistently predicts both the mean and variance of within-population TE load. This model of TE population genetics is then interrogated in order to elucidate those evolutionary genetic mechanisms that have the greatest influence on the population variance of TE load. In particular, moment-based analyses of time-dependent and equilibrium population distributions of TE load reveal that overdispersion is likely to arise via copy-and-paste proliferation dynamics, especially when the elementary processes of proliferation and excision are first-order and balanced. Parameter studies and analytic work confirm this result and further suggest that overdispersed population distributions of TE abundance are probably not a consequence of purifying selection on total element load.

### Classical population genetics of TEs

A good starting point for discussing the evolutionary dynamics of TEs is an influential paper by Charlesworth and Charlesworth (1983) and subsequent work with a similar modeling approach (Brookfield and Badge, 1997; Charlesworth and Charlesworth, 2010; Deceliere, 2004; Le Rouzic and Deceliere, 2005). This classical theory represents a chromosome as a finite set of *m* available insertion sites (loci) per haploid genome, each of which can either be occupied by a transposable element (or not). For a single family of TEs, the state of an infinite diploid population at a given chromosomal site *i*, for *i* = 1, 2,…, *m*, is described by its frequency, *x_i_*, where 0 ≤ *x_i_* ≤ 1. Assuming insertion sites exhibit no linkage disequilibrium, the set of frequencies, 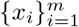, describes the state of the population. The mean copy number of TEs per individual is 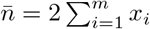, where the factor of 2 accounts for diploidy.

The evolutionarily neutral version of the classical theory includes two processes affecting TE load (gain and loss). Gain of TEs is represented by a proliferation rate (per individual per element per generation) in the germ line of an individual with *n* elements. This proliferation rate, denoted *u_n_*, is typically assumed to be a decreasing function of TE load (*du_n_/dn* < 0). Loss of TEs is represented by a first-order excision rate constant (per individual per element per generation) denoted by *ν*. The change (per generation) in the mean TE copy number per individual is thus

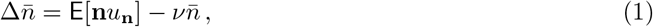

where **n** is the diploid TE load of a randomly sampled individual, the expected value is taken over individuals in the population, and 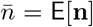 is the population mean of TE copy number. Expanding Eq. 1 around the mean TE load gives the following second-order approximation,

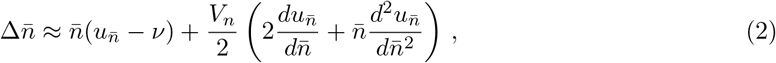

where *V_n_* denotes the population variance in TE copy number (Charlesworth and Charlesworth, 1983). If the higher order terms that scale the population variance are negligible, the change in mean TE copy number per generation is 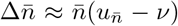. For this neutral model of TE population dynamics, one concludes that 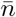 will approach a (stable) equilibrium value satisfying 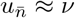 provided 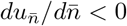.

To extend this model of TE population genetics to include the effect of natural selection, it is customary to assume a viability function, *w_n_*, that is a decreasing function of genome-wide TE load (*dw_n_/dn* < 0). Approximating the mean fitness of the population (E[*w*_n_]) by the fitness of an individual with an average number of copies 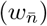, Eq. 2 can be extended to include the effect of selection on TE load (Charlesworth and Charlesworth, 2010),

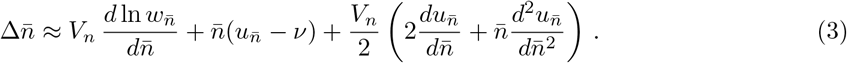

As a specific example, consider the proliferation rate function *u_n_* = *ξ*_0_/*n* with *ξ*_0_ > 0 and the selection function *w_n_* = *e*^−*γn*^ for *γ* > 0 (viability is a decreasing function of TE copy number). Because *du_n_/dn* = −*ξ*_0_/*n*^2^ and *d*^2^*u_n_/dn*^2^ = 2*ξ*_0_/*n*^3^, the higher order terms involving derivatives of *u_n_* evaluate to zero. Consequently, Eq. 3 becomes

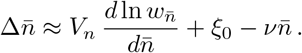

Substituting 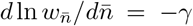 and setting 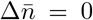 gives 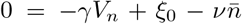. Solving for the equilibrium mean TE load gives,

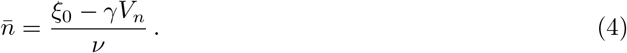

This result is biologically meaningful for *ξ*_0_ > *γV_n_*. As expected, the equilibrium TE load is an increasing function of the proliferation rate constant, *ξ*_0_, and a decreasing function of the excision rate constant, *ν*. Furthermore, stronger selection against TE load (greater *γ*) decreases the mean value of the equilibrium TE load in the population.

### Population variance in the classical model

Analysis of the classical model of TE population genetics proceeds in an *ad hoc* manner by making further assumptions regarding the population variance (*V_n_* is a parameter in Eqs. 2–4). For example, Charlesworth and Charlesworth (1983) assume the population variance takes the form

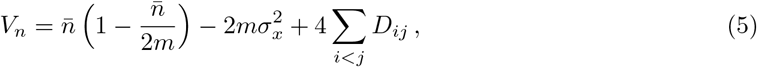

where *D* is a matrix of linkage disequilibrium coefficients (Bulmer, 1980), and 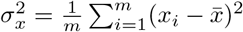 is the variance in element frequency across loci (see Supplemental Material, Section S1). If one further assumes that linkage effects are small enough to be ignored, then

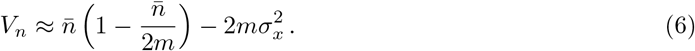

Charlesworth and Charlesworth (1983) argue that for a large enough population, one expects the variance in element frequency across loci to be eventually become negligable, 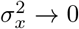 and, consequently, the equilibrium population variance of TE load should approach that of a binomial distribution,

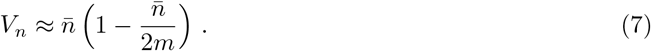

In that case, assuming occupiable loci are not limiting 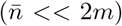, the population variance will be well-approximately by the mean 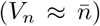. Substituting this value into Eq. 4, the classical model indicates that the equilibrium TE load will be

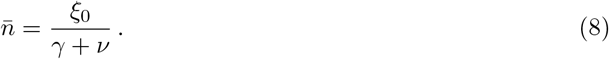

As in Eq. 4, the equilibrium TE load is an increasing function of the proliferation rate constant (*ξ*_0_), and a decreasing function of both the excision rate constant (*ν*) and the strength of selection against TE load (*γ*).

The classical model (Eqs. 3–8) has informed expectations regarding the population genetics of TEs for several decades. For example, an extension of this classical theory predicts that in a finite population of effective size *N_e_*, the the stationary distribution of TE frequency (*x*) will take the form *ρ*(*x*) ∝ *x*^*a*–1^(1 – *x*)^*b*–1^ where 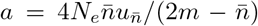 and 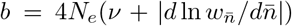 (Le Rouzic and Deceliere, 2005). For *u_n_* = *ξ*_0_/*n*, *w_n_* = *e*^−*γn*^, and 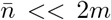, this gives *a* = 4*N_e_ξ*_0_ and *b* = 4*N_e_*(*ν*+*γ*). On the other hand, the classical approach to modeling TE population genetics has obvious limitations. For one thing, the derivation and analysis of the classical model makes assumptions about the population variance (*V_n_* in Eqs. 5–7) that may not be consistent with experimental observations (see Results). Furthermore, the population variance of TE load ought to be an emergent property of a model constructed for the purpose of understanding the population genetics of TEs, rather than a modeling assumption imposed upon a preexisting framework (Eq. 7).

The remainder of this paper addresses these two issues in detail. We begin by presenting empirical evidence that population variance of TEs is neither binomial nor well-approximated by the mean. This motivates the presentation of an alternative population genetic framework that simultaneously and self-consistently predicts both the population variance and the mean TE load. This model of TE population genetics is then interrogated in order to elucidate those evolutionary genetic mechanisms that have the greatest influence on the population variance of TE load.

## Results

### Dispersion of TE loads in the classical model

In the classical modeling of TE population genetics discussed above, analytical results are obtained by assuming a randomly mating population with a binomial distribution of TE loads,

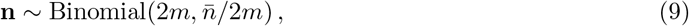

with mean 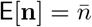 and variance 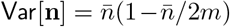 (Eqs. 5–7). A simple measure of the variability of TE load within a population is the *index of dispersion* (Fano factor) given by

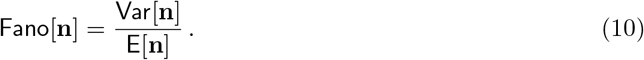

Substituting the mean and variance of the binomial distribution into Eq. 10, it is apparent that the classical model of TE population genetics predicts (i.e., assumes) a Fano factor that is less than one,

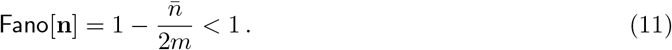

In fact, when the number of sites occupied by TEs is small compared to the total number of occupiable loci (m → ∞ with 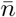 fixed), the Fano factor approaches one from below (Fano[**n**] → 1). In this limit, the binomial distribution of Eq. 9 is well-approximated by 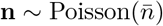. If it were the case that the TE load within a population were Poisson distributed, then the mean and variance of TE load would be equal 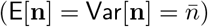 and the index of dispersion would be Fano[**n**] = 1. With our expectations set by this prediction of the classical model of TE population genetics, empirical observations of a Fano factor greater than one (Fano[**n**] > 1) would indicate *overdispersion* of TE load within a population.

### Overdispersion of empirical TE counts

Figs. 1 and 2 present analyses of two data sets, both of which demonstrate that the variance of TE load in experimentally studied populations can be far greater than would be predicted by classical models of TE population genetics. The first data set (analyzed in Fig. 1) consists of whole-genome sequence data from 164 lines of *Mimulus guttatus* derived from a naturally occurring population (hundreds of thousands of individuals) in Iron Mountain, Oregon, USA (Troth et al., 2018). To estimate TE copy numbers, we compared the coverage of each TE to the average coverage of single copy genes (see Appendix for details). Although *Mimulus guttatus* is our primary interest, we also analyzed, for comparison, a data set (Fig. 2) that consists of genomic DNA sequencing data from 131 lines of *Drosophila melanogaster* (derived from a large population in Raleigh, North Carolina) obtained from the Drosophila Genetic Reference Panel (Cridland et al., 2013, Supplementary Material S5).

**Figure 1:**
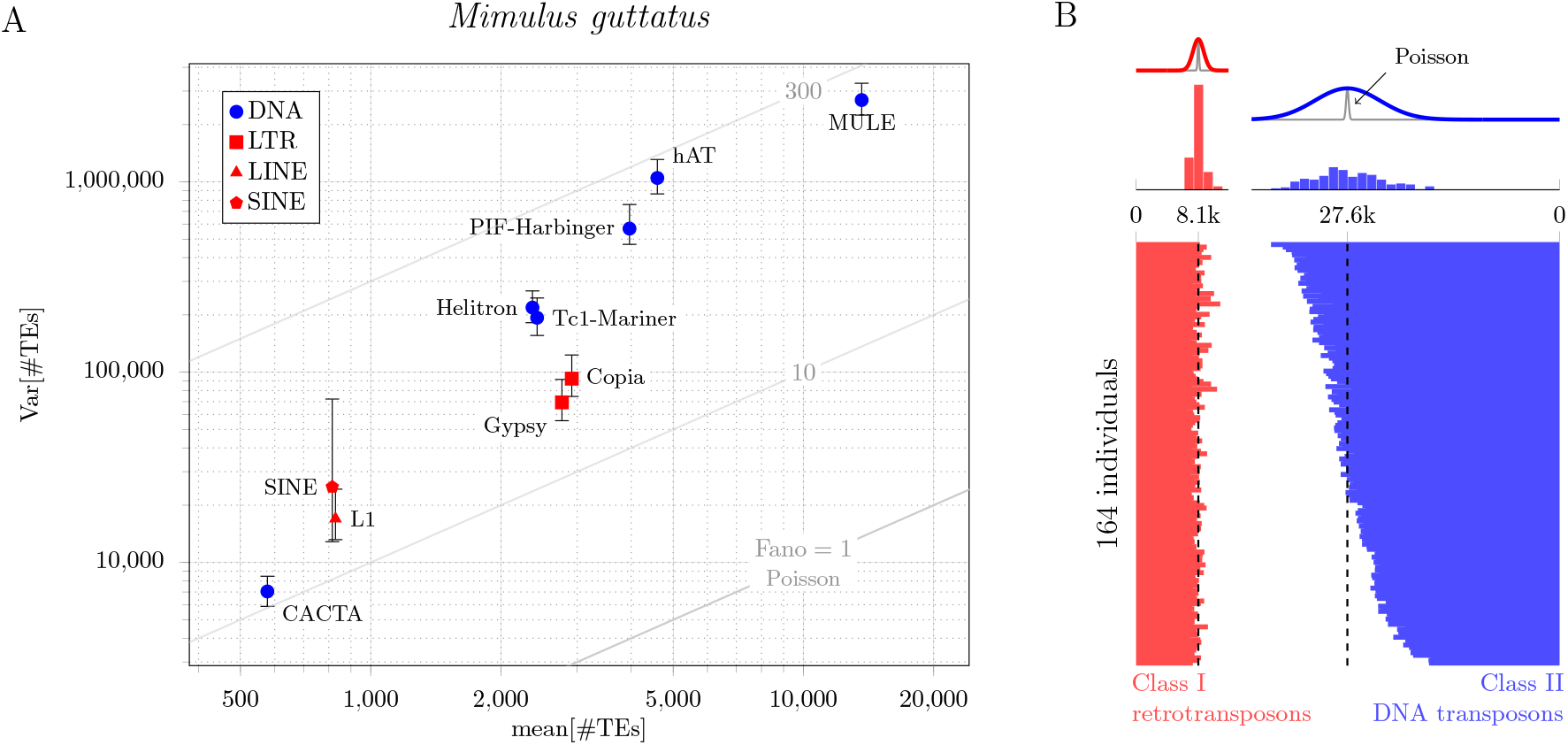
A: Mean-variance plot of TE copy number in a *M. guttatus* population compared to theoretical expectation (Poisson line). For each of ten different families of TEs, the index of dispersion is in the range 10 < Fano[**n**] < 300. For each family, the vertical bars show 95% bootstrap confidence interval of the population variance of TE load. B: Estimated TE copy number for 164 *M. guttatus* individuals. TE counts are separated by class (red, Class I, retrotransposon; blue, Class II, DNA transposon). The variability in TE load can be observed in the counts from individuals (bottom) as well as histograms (top). The overdispersion in TE load is apparent in the deviation of the observed counts (red and blue histograms) from the corresponding Poisson distributions (gray lines). The vertical dashed lines show the population mean of TE load.

**Figure 2:**
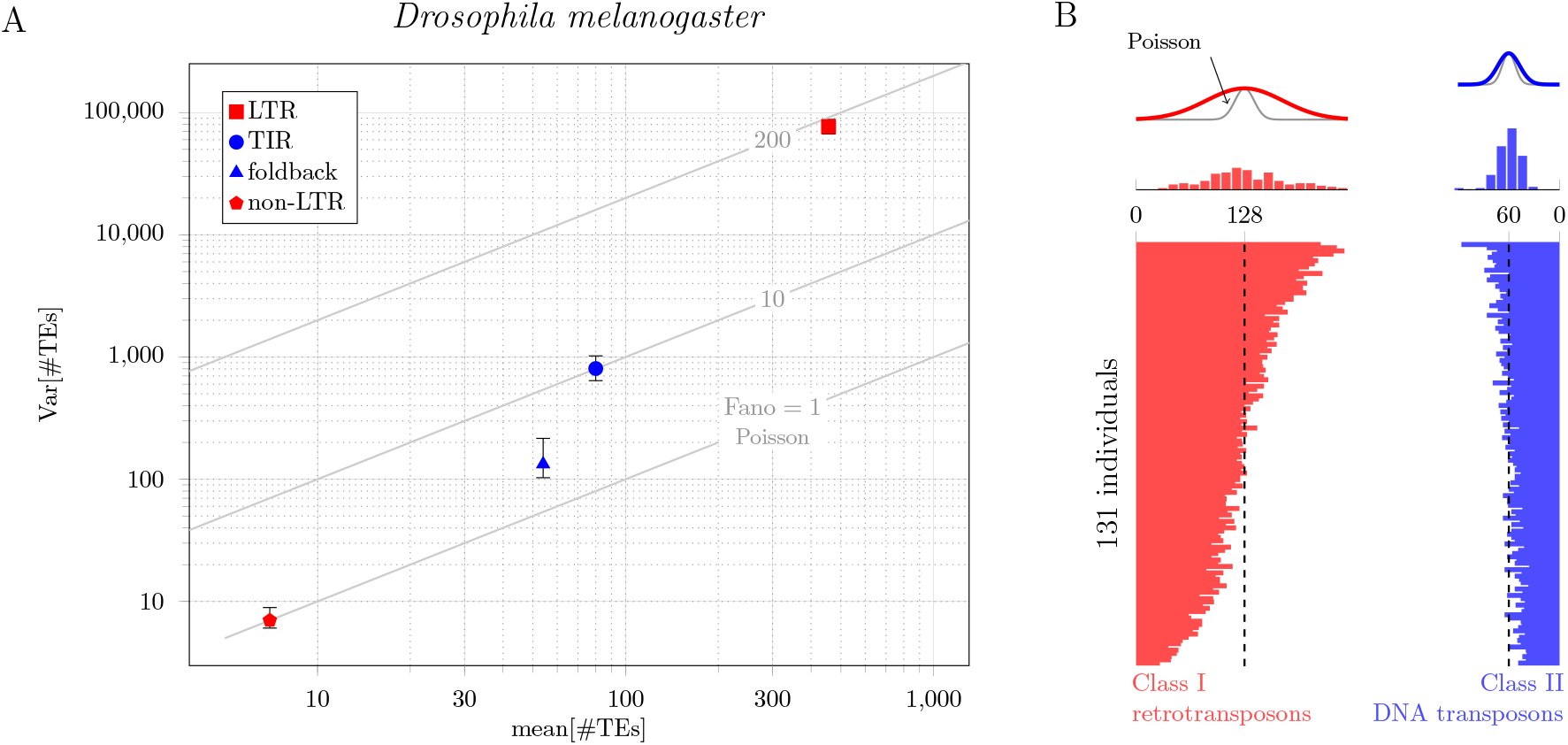
A: Mean-variance plot of TE copy number in *D. melanogaster* compared to theoretical expectation (Poisson line). For each of four different families of TEs, the index of dispersion is in the range 1 < Fano[**n**] < 200. Vertical bars show 95% bootstrap confidence intervals. B: Estimated TE copy number for 131 *D. melanogaster* individuals. TE counts are separated by class (red, Class I, retrotransposon; blue, Class II, DNA transposon).

**Figure 3:**
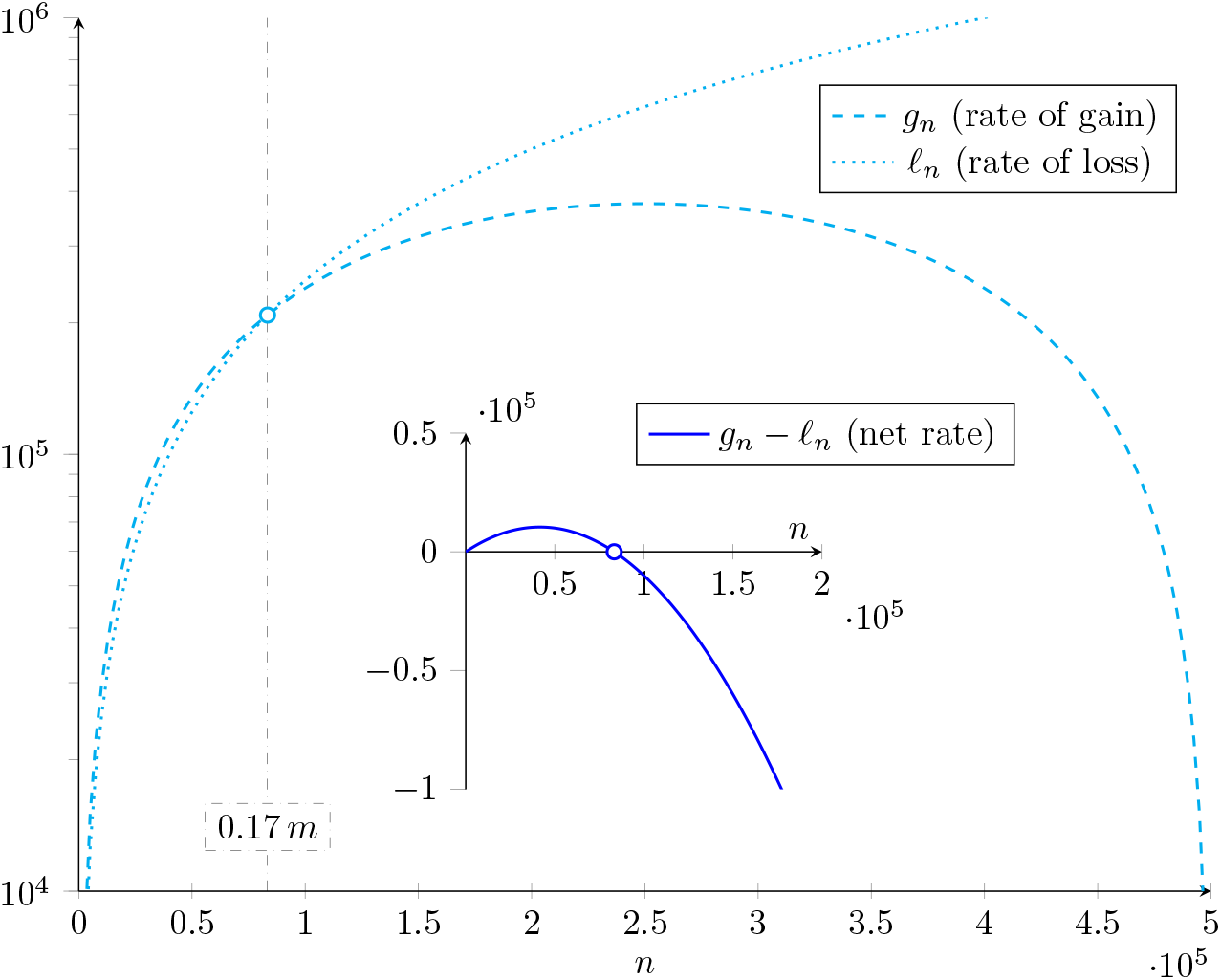
Example rates of transposable element gain (*g_n_*, dashed) and loss (*ℓ_n_*, dotted) are shown cyan. These functions of TE load given by Eqs. 20–21 intersect (balance) when 17% of the insertion sites are occupied (gray dot-dashed line). Parameters: *η*_0_ = 20, *η* = 3, *ν* = 2.5, *m* = 5 × 10^5^. The net rate of change (blue curve) is zero for a TE load of *n* = 8.3 × 10^4^ (open circle), which is on the order of that found for transposons (Class II elements) in *Mimulus* (e.g., LINE and LTR, see Fig. 1B left).

Comparison of the marker locations and histograms in Figs. 1 and 2 with the gray lines labelled Poisson shows that in both species, *Mimulus guttatus* and *Drosophila melanogaster*, the population distribution of TE load is *overdispersed* (the variance of TE load is greater than the mean TE load). In *D. melanogaster*, this overdispersion is greater for Class I TEs (retrotransposons) with an RNA intermediate than Class II TEs (DNA transposons). (Fig. 2B). The corresponding Fano factors (Eq. 10) are 16 and 2.7, respectively; these values indicate overdispersion (see Table 1). Fig. 1A shows that overdispersion of TE load is even more pronounced in *M. guttatus*. In this case, the Fano factors are 61 for Class I TEs (red symbols: LINE, SINE and LTR), and 646 for Class II TEs (blue symbols: CACTA, Helitron, Tcl-Mariner, PIF-Harbinger, hAT and MULE). Fig. 1B shows the estimated number of Class I and II TEs in each of the 164 lines of *M. guttatus* (horizontal bar graph). In both cases, the variance (illustrated by the width of red and blue histograms) is far greater than the variance that would be consistent with classical model of TE population genetics (gray curves). Taken together, Figs. 1 and 2 show that in both species, *Mimulus guttatus* and *Drosophila melanogaster*, and for both classes of TEs (retrotransposons and DNA transposons), the distribution of TE load within the studied population is highly *overdispersed*.

**Table 1:** Empirically observed mean, variance, and index of dispersion (Fano factor) of the population distribution of TE load in 164 *M. guttatus* and 131 *D. melanogaster* individuals (cf. Figs. 1 and 2). Class I elements (retrotransposons) proliferate in a staged manner that involves an RNA intermediate, while Class II elements (DNA transposons) do not utilize an RNA intermediate (see, e.g., Ch. 9 of Graur (2016) for review).

| Species | TE Class | $E[\mathbf{n}]$ | $\text{Var}[\mathbf{n}]$ | $\text{Fano}[\mathbf{n}]$ |
| --- | --- | --- | --- | --- |
| *M. guttatus* | I | 8,082 | $4.9 \times 10^5$ | 61 |
| | II | 27,559 | $1.8 \times 10^7$ | 646 |
| *D. melanogaster* | I | 128 | 2,053 | 16 |
|  | II | 60 | 164 | 2.7 |

### Overdispersion is not explained by distinct TE families

The overdispersion documented in Figs. 1 and 2 cannot be explained away as a simple consequence of the heterogeneity of the properties of distinct TE types. Consider two families of TEs with loads across individuals in the population given by the random variables **x**_1_ and **x**_2_. Denoting the mean TE loads of these families by 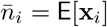, the corresponding Fano factors are 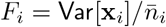. If these two families were not distinguished, the observed mean load would be given by a composite count, **x** = **x**_1_ + **x**_2_, with mean 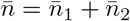 and variance,

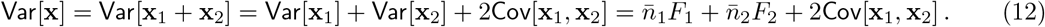

Substituting 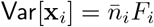 and dividing by 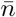 gives an expression for the composite index of dispersion,

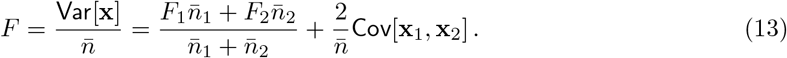

Assuming that the within-population loads for the two families of TEs are independent, the covariance will be zero (Cov[**x**_1_, **x**_2_] = 0). In that case, the composite Fano factor is a weighted average of Fano factors for each family,

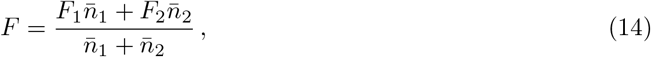

which takes values in the range min(*F*_1_, *F*_2_) ≤ *F* ≤ max(*F*_1_, *F*_2_). A similar argument allows us to conclude that for TE families with independent proliferation and excision dynamics, the dispersion of TE load that results when families are not distinguished is always *less* than the overdispersion of at least one of the TE families (see Section S2 of the Supplemental Materials for further discussion).

### Master equation for TE population dynamics

Our modeling aims to clarify the observed overdispersion of TE load in *Mimulus guttatus* and *Drosophila melanogaster*, following classical TE population genetics, but with a few important modifications. Because the variance in TE load is not the result of heterogeneity in TE types (see above), our analysis will focus on a single TE family.

Let *p_n_*(*t*) denote the probability that a randomly sampled haploid genome (gamete) has a TE count of *n* at time *t*. Prior to considerations of selection, our neutral model of TE population dynamics will take the form of a skip-free birth-death process with gain and loss rates denoted *g_n_* and *ℓ_n_*. The discrete state space for haploid TE load is **n** ∈ {0, 1, 2,…, *m*} and the state-transition diagram of the stochastic process is

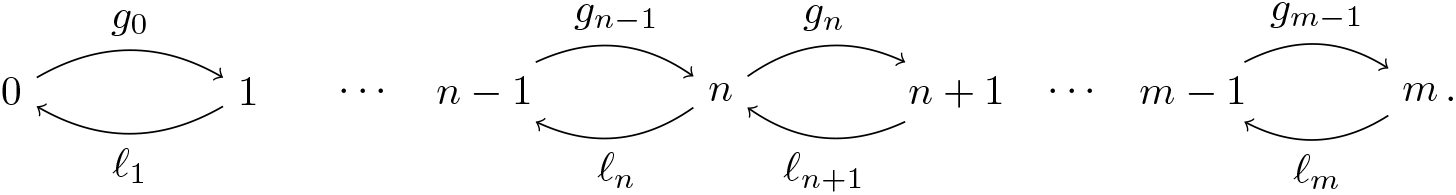

The master equation for this stochastic process is the following system of *m* +1 differential equations,

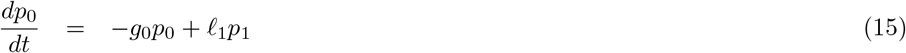

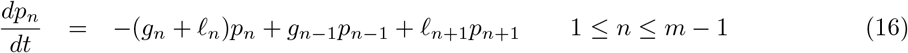

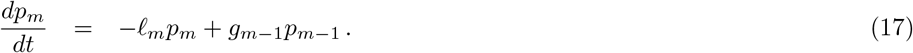

The expected value of TE load of a randomly sampled diploid genotype is

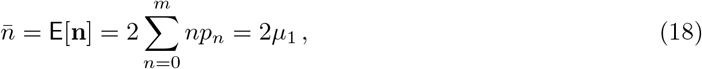

where 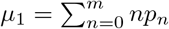 is the mean TE load of a randomly sampled haploid gamete. By differentiating Eq. 18 to obtain

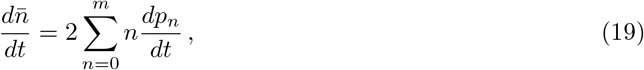

and substituting Eqs. 15–17, the master equation formulation can be shown to be consistent with the classical model (see Section S3).

### Master equation predicts the variance of TE load

One feature of the master equation formulation (Eqs. 15–17) is that the dynamics of the population variance of TE load are an emergent property of the model. To illustrate, let us assume that TE excision occurs with first order rate constant, then the loss rate as a function of *n* is

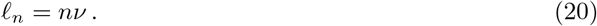

Let us further assume that the rate of gain for a single family of TEs takes the form

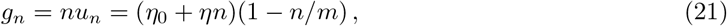

where *η* is the copy-and-paste rate per transposon (a first-order rate constant characterizing proliferation of TEs), *η*_0_ is a zeroth order rate constant, *n* is the TE copy number, and *m* is the number of occupiable loci (in a haploid gamete). Substituting these constitutive relations for the loss and grain rates, *ℓ_n_* and *g_n_*, into Eqs. 15–17 gives

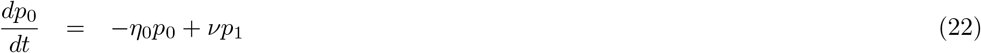

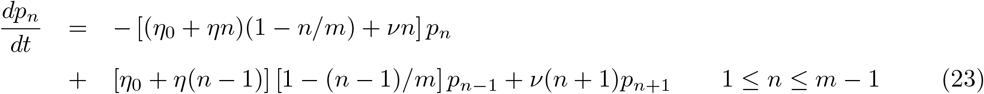

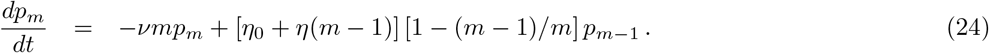

Fig. 4 shows representative numerical solutions of this master equation for the population dynamics of TE load. When the copy-and-paste rate constant is zero (*η* = 0) and occupiable loci are not limiting 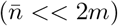, the stationary probability distribution is well-approximated by a Poisson distribution with 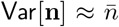 and Fano[**n**] ≈ 1 (blue histograms). For both *Mimulus*- and *Drosophila*-like parameters, no overdispersion is observed when *η* = 0. These results should be compared to the green and red histograms, for which the copy-and-pate rate is nonzero (see caption for parameters). Notably, an increase in the copy-and-paste rate leads to significant overdispersion of the TE load for both simulated populations (Fano[**n**] ranging from 1 to 100).

**Figure 4:**
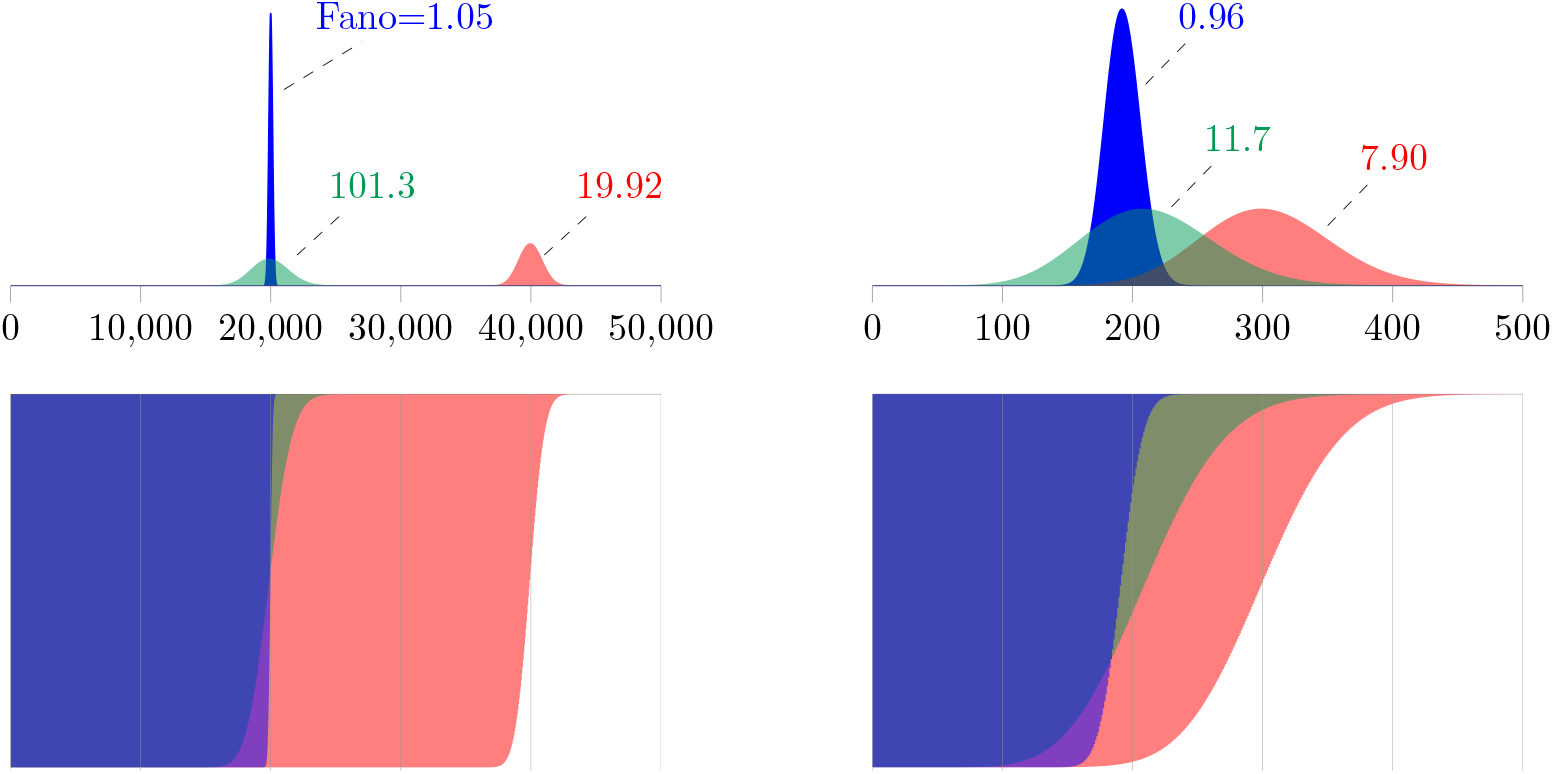
Top: Stationary population distributions of TE load in haploid genomes (gametes) calculated using the evolutionarily neutral master equation model (Eqs. 22–24). For mean loads similar to *Mimulus* (left) and *Drosophila* (right), no overdispersion is observed in simulations absent copy-and-paste transposition (*η* = 0, Fano[**n**] ≈ 1). Green and red histograms show overdispersed population distributions of TE load that are obtained when copy-and-paste transposition is included. *Mimulus* parameters: *ν* = 0.1, *m* = 10^9^; *η*_0_, *η* = 2000, 0 (blue), 200, 0.095 (red), 20, 0.099 (green). *Drosophila* parameters: *ν* = 0.1, *m* = 5000; *η*_0_, η = 20, 0 (blue), 1, 0.1 (green), 2, 0.1 (red). See Section S6 for numerical methods.

### Moment equations for mean and variance of TE load

The previous section showed that the evolutionarily neutral master equation model provides information about the population variance of TE load that is unavailable in classical theory. Because this realism comes at the expense of a more complex model formulation (Eqs. 22–24 compared to Eq. 2), we derived ordinary differential equations (ODEs) that summarize the dynamics of the mean and variance of the population distribution of diploid TE loads predicted by the master equation. Section S3 of the Supplementary Material shows that the mean and variance of TE load solve the following ODEs,

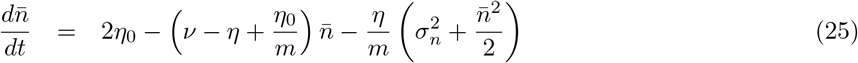

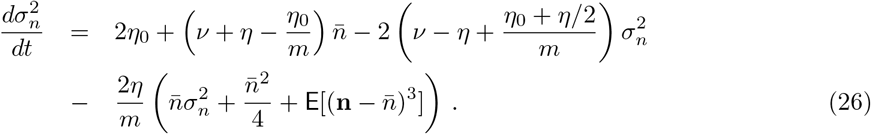

The term 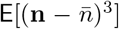 that appears in Eq. 26 is the third central moment of the within-population diploid TE load. Analysis of this system of ODEs and the third central moment is provided below.

If number of occupiable loci are not limiting 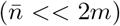, we may take the limit of Eqs. 25–26 as *m* → ∞ to obtain simpler equations for the mean and variance,

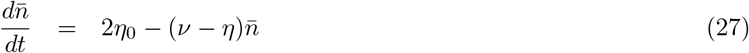

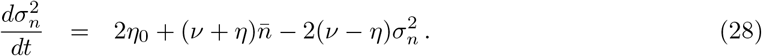

This reduced system of ODEs is linear and, for large *m*, the equation for the variance (Eq. 28) does not depend on the third central moment. The steady-state solution of Eqs. 27–28 is given by

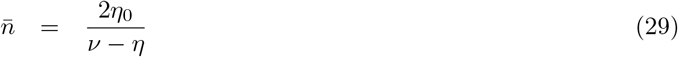

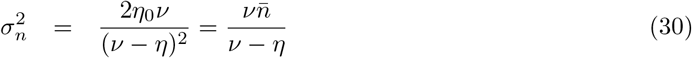

is physical provided *ν* > *η*, that is, when *m* is large, the rate of excision *ν* must be greater than the copy-and-past rate constant *η* for a biologically meaningful solution with 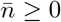 (mean TE load must be non-negative). This steady state is stable because the Jacobian of Eqs. 27–28, given by the 2 × 2 matrix with entries *J*_11_ = −(*ν* – *η*), *J*_12_ = 0, *J*_21_ = *ν* + *η*, *J*_22_ = −2(*ν* – *η*), has real valued eigenvalues λ_+_ = −(*ν* – *η*) < 0 and λ_−_ = 2λ_+_ < 0.

The values for the steady-state mean and variance of TE load given by Eqs. 29–30 correspond to the following index of dispersion,

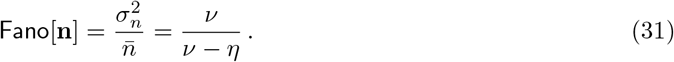

Notably, the condition for a stable steady state (ν > η) implies an index of dispersion greater than unity (Fano[**n**] > 1) for any nonzero copy-and-paste rate constant (*η* > 0). For this reason, we conclude that *copy-and-paste proliferation dynamics will result in an overdispersed steady-state population distribution of TE loads provided the number of occupiable loci are not limiting* 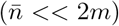. Further analysis of the moment equations (Eqs. 25–26) shows that overdispersion will not occur in the absence of copy-and-paste dynamics (see the *η* = 0 case in Table 2).

**Table 2:**
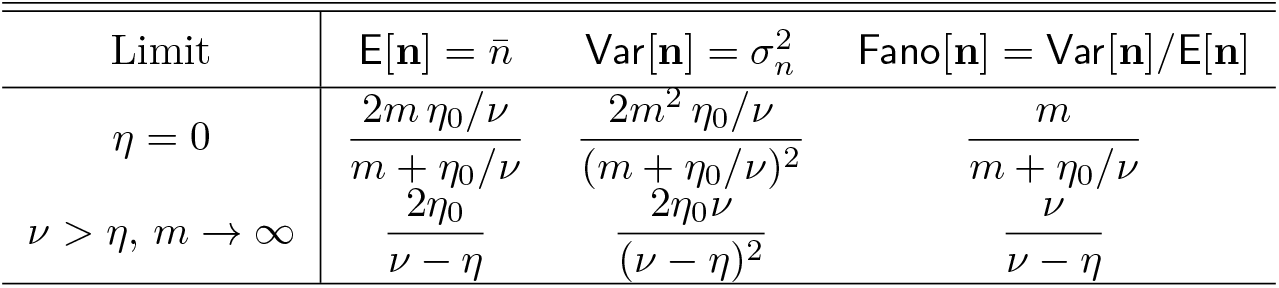
The evolutionarily neutral moment equations (Eqs. 25–26) for the mean and variance of TE load make predictions in various limits (see Sections S3.2–S3.3).

This preliminary analysis of an evolutionarily neutral master equation for TE proliferation (Eqs. 22–24) indicates that *a nonzero copy-and-paste rate may lead to an overdispersed population distribution of TE load* (Eq. 31). That is, copy-and-paste TE dynamics is one possible explanation for our empirical observations of overdispersed TE counts (Figs. 1 and 2). Furthermore, this analysis predicts that a large index of dispersion may be a consequence of balanced dynamics of TE gain and loss (i.e., Fano[**n**] → ∞ as *ν* decreases to *η* in Eq. 31). While the divergence in the analytical result is an artifact of taking the *m* → ∞ limit, a parameter study of the master equation model (Fig. 5) confirms that overdispersion is most pronounced (Fano[**n**] maximized) when *m* is large and the dynamics of TE gain and loss are approximately balanced (*η* ≈ *ν*).

**Figure 5:**
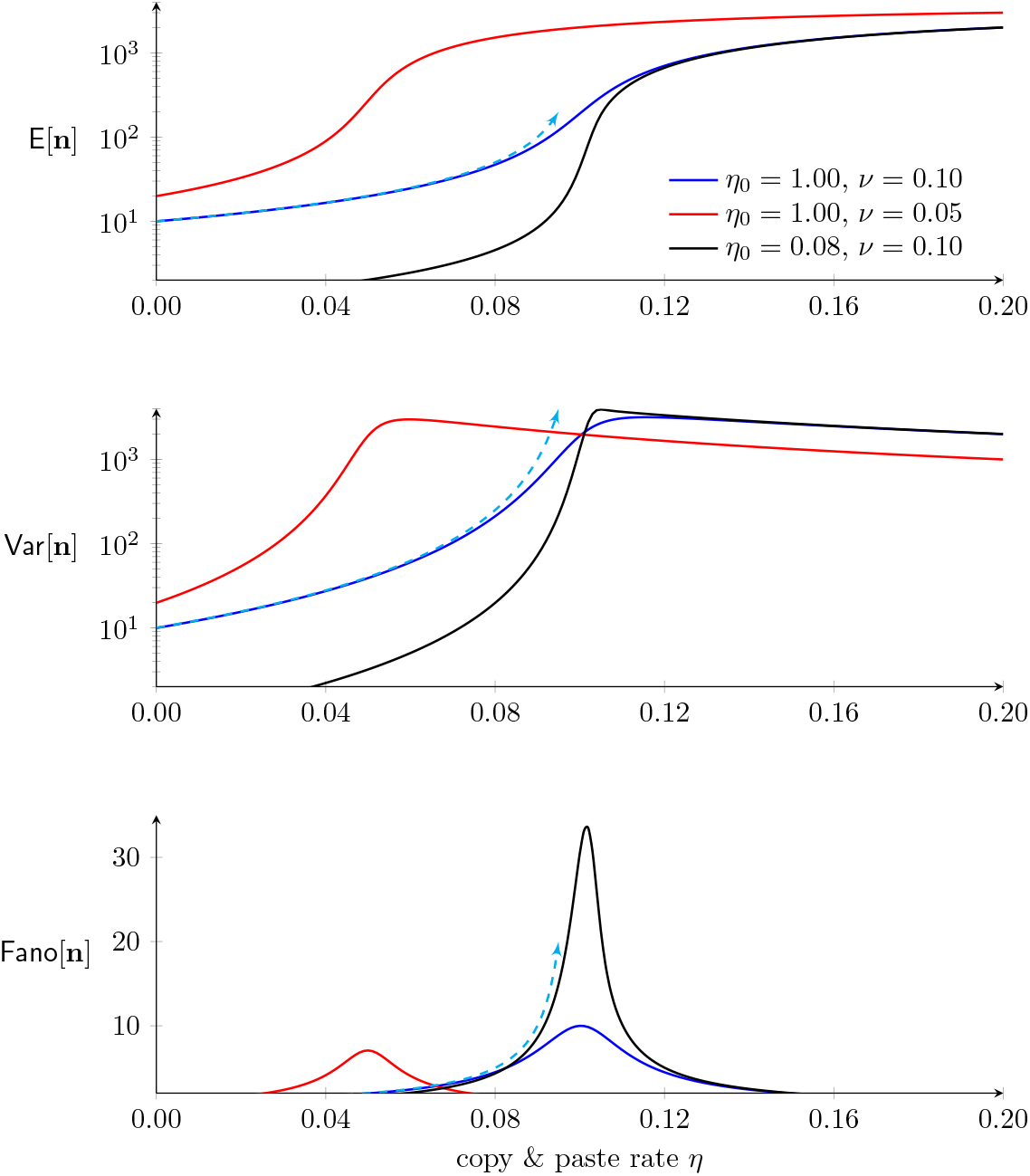
Parameter studies of the neutral master equation model showing the mean 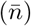, variance 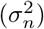, and index of dispersion (Fano[**n**]) of within-population TE load as a function of the copy-and-paste rate constant (*η*). Parameters: *m* = 4 × 10^3^ and as in legend. Cyan curves indicate analytical approximations using *ν* = 0.1 that are valid in the limit as *m* → ∞ (see Table 2). These approximations are most accurate for small *η*/*ν* and diverge as *η* approaches *ν* from below (cyan arrowheads). These calculations were accelerated using a Fokker-Planck approximation to Eqs. 22–24 (see Section S6).

### Influence of selection on overdispersion

To investigate the effect of purifying selection on the population variance of TE load, we assume a selection coefficient (*w_n_*) that depends on total diploid TE load (*n*) with *dw_n_/dn* < 0 (higher load is less viable). For concreteness, let

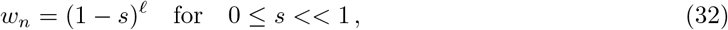

where *s* is the strength of selection against TE load. When the neutral model (Eqs. 22–24) is modified to include selection, the master equation becomes

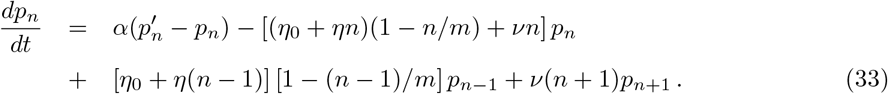

for 1 ≤ *n* ≤ *m*. The first term in this expression represents each load probability *p_n_* relaxing to a target probability 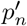 given by

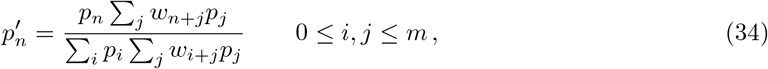

where *w*_*i*+*j*_ = (1 – s)^*i*+*j*^. The equations for for *dp*_0_/*dt* and *dp_m_/dt* have fewer gain/loss terms than Eq. 33, but are analogous (cf. Eqs. 22 and 24). The parameter *α* that occurs in Eq. 33 is the inverse of the generation time. The quantity 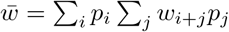 is the mean fitness under the assumption of random mating (Gillespie, 2004).

Fig. 6 shows steady-state distributions of haploid (top row) and diploid (bottom) TE loads calculated using Eq. 33 both with and without of selection on diploid load. As expected, for both *Mimulus*- and *Drosophila*-like mean TE loads, the effect of weak selection (red and green histograms) is to decrease the TE load in the population as compared to the neutral model (blue histograms). This decrease in mean TE load occurs for a wide range of generation times (1/*α*) and selection coefficients (*s*). More important (and less obvious) is the impact of selection on the variance of TE load and overdispersion. Using *Drosophila* parameters, Fig. 6 (top right) shows an example simulation (green histogram) in which selection leads to increased dispersion (the Fano factor increases from 1 to 8.66). However, in a second case (red histogram), selection increases the index of dispersion only slightly (to a Fano factor of 1.06). Notably, in three representative simulations using *Mimulus* parameters, selection does not increase the dispersion of TE load (Fig. 6, left). This observation is consistent with the moment-based analysis presented in the following section.

**Figure 6:**
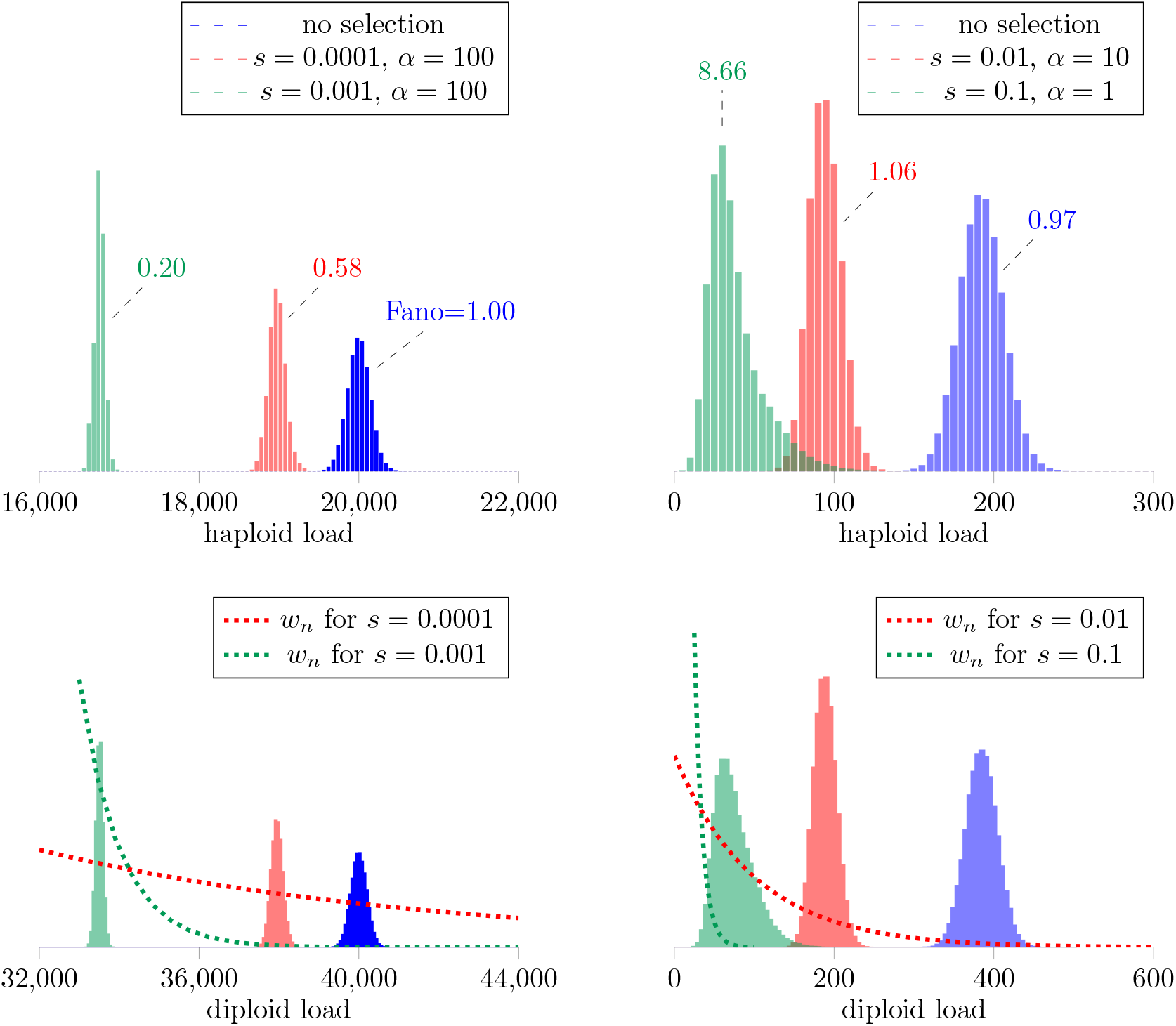
Stationary population distributions of TE abundance with and without selection given by numerical solution of Fokker-Planck equation associated to the master equations (Eqs. 33–34). Parameters: selection coefficient s as in legends. *Mimulus*: *ν* = 0.1, *m* = 10^9^; *η*_0_, *η* = 2000, 0 (blue), 200, 0.095 (red), 20, 0.099 (green). *Drosophila*: *ν* = 0.1, *m* = 5000; *η*_0_, *η* = 20, 0 (blue), 1, 0.1 (green), 2, 0.1 (red).

### Moment equations with selection

For a deeper understanding of the impact of selection on the distribution of TE load in a population, one may begin with Eqs. 33–34 and derive the dynamics of the mean and variance of TE load under the action of simple selection functions. For example, in the limit of weak selection 0 < s ≪ 1, Eq. 32 is well-approximated by *w_n_* = 1 – *sn*. In this case, as derived in Section S4, the dynamics of the mean and variance of TE load solve

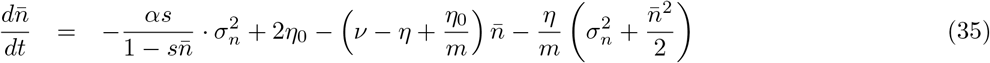

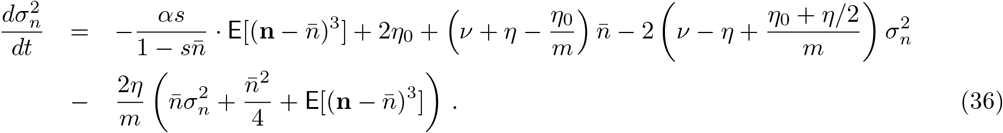

These ODEs may be compared to the moment equations for the neutral model (Eqs. 27–28). As expected, the influence of selection on the mean TE load is proportional to the population variance (through the factor 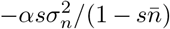 in Eq. 35). Similarly, the influence of selection on the population variance is proportional to the third central moment of the diploid load (through the factor 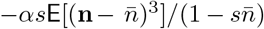 in Eq. 36). In both cases, the quantity 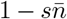 is the mean fitness of the population, i.e., 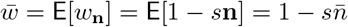.

Under the assumption that the mean TE load is much smaller than the number of loci 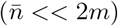, we may simplify the moment equations with selection (Eqs. 35–36) by taking the limit *m* → ∞ to obtain

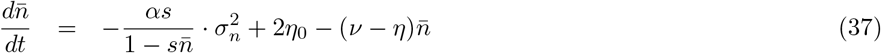

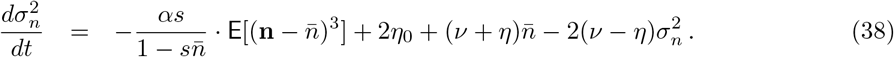

Setting the left side of Eq. 37 to zero, we observe that the steady-state mean and variance are related as follows,

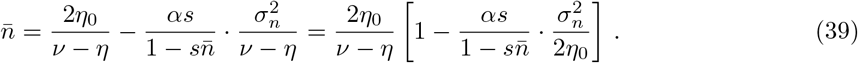

Comparing this expression to Eq. 29, noting that the variance is nonnegative 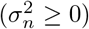, we see that the effect of weak selection (0 < *s* ≪ 1) is to decrease the mean TE load in the population as compared to the neutral model (as expected). Similar analysis of Eq. 38 shows how selection may impact the variance of TE load and, consequently, the index of overdispersion. Setting the left side of Eq. 38 to zero and solving for the steady-state variance, gives

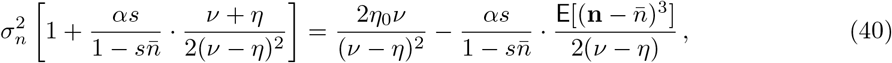

where the first term on the right side, 2*η*_0_*ν*/(*ν* – *η*)^2^, is the variance of TE load in the absence of selection. Eq. 40 shows that the effect of selection is to either decrease or increase the population variance of TE load, depending on the sign of the third central moment 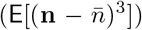, which is consistent with the master equation simulations shown in Fig. 6.

### Moment closure and the 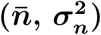 phase plane

In their current form, the moment equations (Eqs. 35–36) are an open system of ODEs, because the equation for the variance 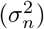 depends on 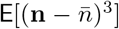, the unknown third central moment. As discussed in Section S5, a moment closure technique that is applicable in this situation assumes that the third central moment of the diploid load is algebraic function of the mean and variance,

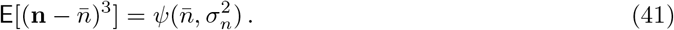

We investigated two possibilities for this function based on the properties of the beta-binomial and negative binomial distributions. The beta-binomial moment closure, derived in Section S5.3, is a complicated expression involving the mean, variance, and number of loci *m*,

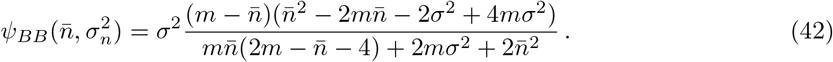

Moment closure motivated by the properties of the negative binomial distribution results in a simpler expression that does not involve the number of loci *m*,

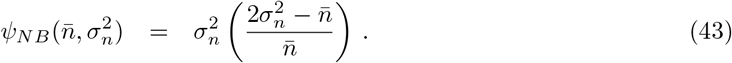

Although the beta-binomial closure (Eq. 42) is arguably a better approximation, in our experience it does not perform markedly better than the negative binomial closure (Eq. 43), as assessed through comparison of moment ODE and master equation simulations. In the analysis that follows, we use the negative binomial closure, motivated by its simplicity and the fact the two expressions coincide when the number of loci are not limiting (to see this, observe that *ψ_BB_* → *ψ_NB_* as *m* → ∞). When the algebraic relationship representing the negative binomial closure (Eq. 43) is substituted into Eqs. 25–26, we obtain the following system of ODEs for the mean and variance of diploid load under the influence of selection:

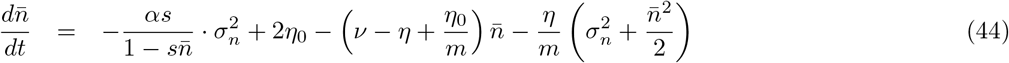

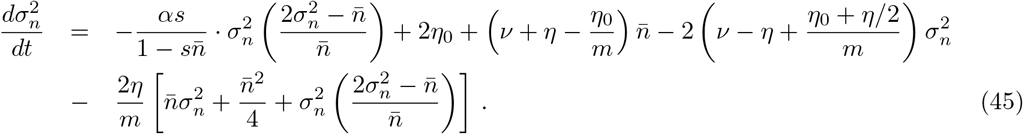

Fig. 7A presents a representative (*n*, 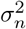) phase plane for the dynamics of the mean and variance of TE load predicted by Eqs. 44–45. The red and green lines are the nullclines for the mean and variance, respectively, with intersection corresponding to the steady state. This calculation uses parameters resulting in a steady-state TE load similar to our empirical observations of *Mimulus guttatus* (counts on the order of 10^5^). This steady state predicted by the moment equations is located far above the broken black line denoting 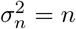 and Fano factor of 1. The blue curves show two solutions that use initial conditions for which the population variance is equal to the mean (solutions were obtained by numerically integrating Eqs. 44–45). Interestingly, these solutions show that dynamics of TE load can include a transient phase in which the index of dispersion is far greater or less than the steady-state value.

**Figure 7:**
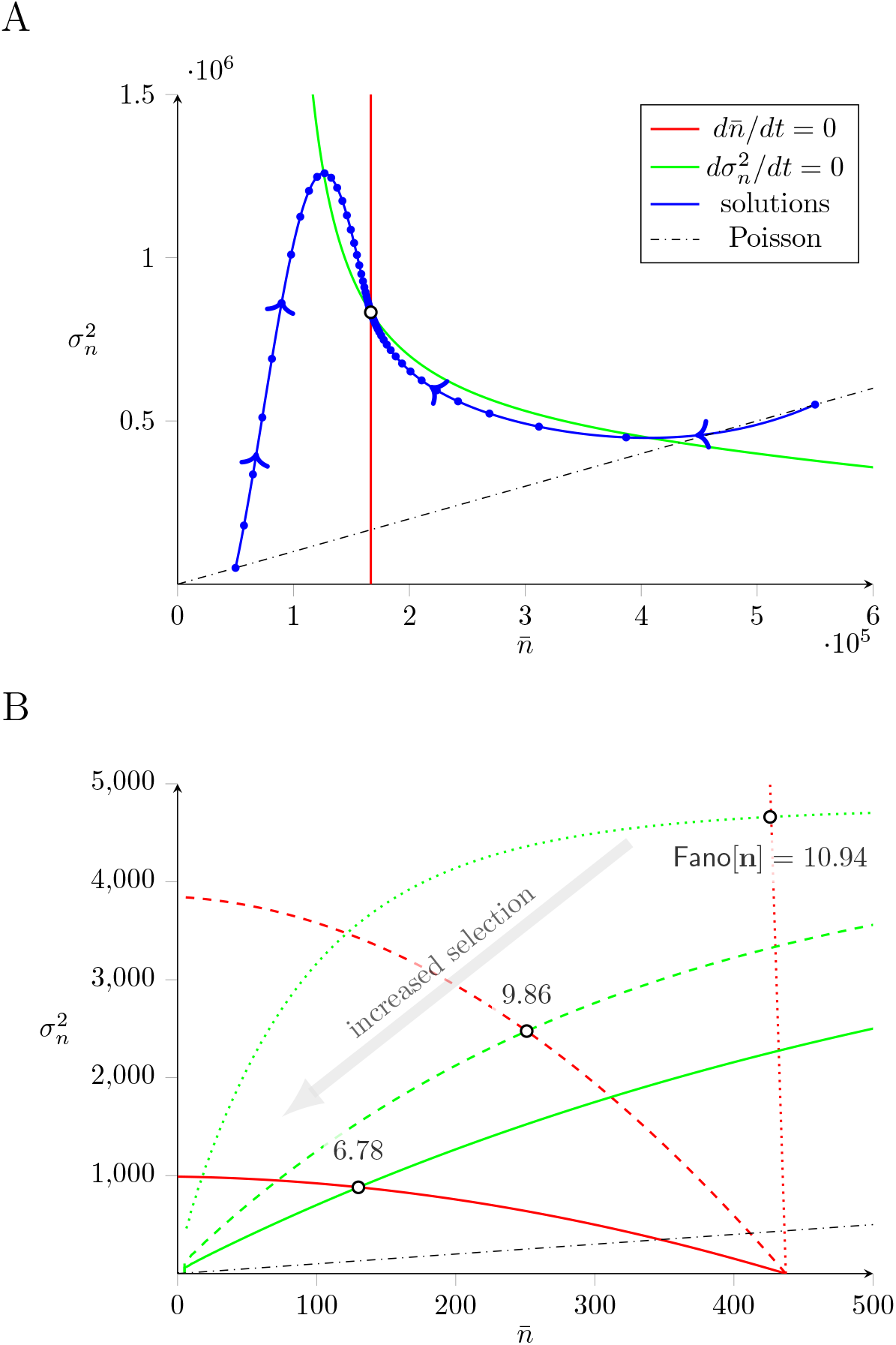
The phase plane for the dynamics of mean (*n*) and variance 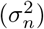 of TE load (Eqs. 44–45). The red and green curves are the nullclines for the mean and variance, respectively, with intersection corresponding to the steady state (open circle). The blue trajectories show the dynamics of equilibration. A: Mean loads similar to *Mimulus*. Parameters: *ν* = 2.5, *m* = 5 × 10^5^, *η*_0_ = 20 and *η* = 3 with no selection (*α* = 0, *s* = 0). B: Mean loads similar to *Drosophila*. Parameters: *ν* = 0.1, *m* = 5000, *η*_0_ = 1 and *η* = 0.1. Dotted nullclines: no selection. Dashed: *s* = 10^-4^, *α* = 5. Solid: *s* = 10^-4^, *α* = 20. Note that increased selection on TE load (gray arrow) decreases the index of dispersion (Fano[**n**]).

Fig. 7B shows how the nullclines for the mean and variance of TE load depend on the strength of selection in three cases with parameters corresopnding to TE loads similar to *Drosophila melanogaster* (counts on the order of 100). As the strength of selection increases, both the mean and variance of TE load decrease, in such a manner that the index of dispersion (Fano[**n**]) decreases.

Although the model obtained by moment closure and the phase plane analysis of Fig. 7 does not assume 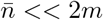, we may consider Eqs. 44–45 in the limit as *m* → ∞,

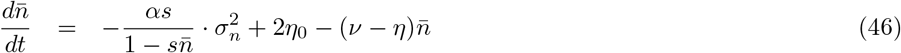

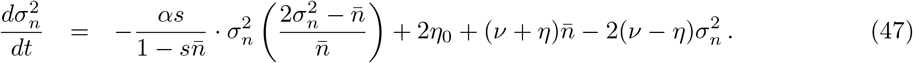

Setting the left sides of Eqs. 44–45 to zero, and assuming weak selection (0 ≤ *s* ≪ 1), we can derive first-order accurate asymptotic expressions for the steady-state mean and variance,

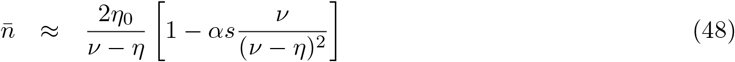

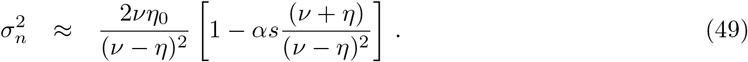

Because *υ*/(*ν* – *η*)^2^ > 0, this expression indicates that weak selection decreases the mean TE load, consistent with our intuition. Similarly, the factor (*ν* + *η*)/(*ν* — *η*)^2^ is positive, so we conclude that weak selection decreases the population variance when *m* is large. As for the index of dispersion, this analysis indicates that under weak selection the Fano factor is

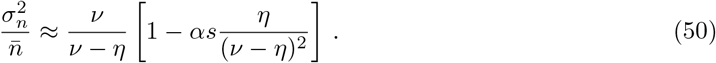

Because *η*/(*ν* – *η*)^2^ is positive any nonzero copy-and-paste rate (*η* > 0), we conclude that the Fano factor is also expected to decrease, because weak selection causes the within-population variance of TE load to decrease more than the mean. This conclusion—i.e., selection on diploid TE load is unlikely to be responsible for overdispersion—is consistent with the numerical parameter studies summarized in Fig. 8 that were enabled by the moment equations with selection (Eqs. 35–36) and beta-binomial moment closure (Eq. 42).

**Figure 8:**
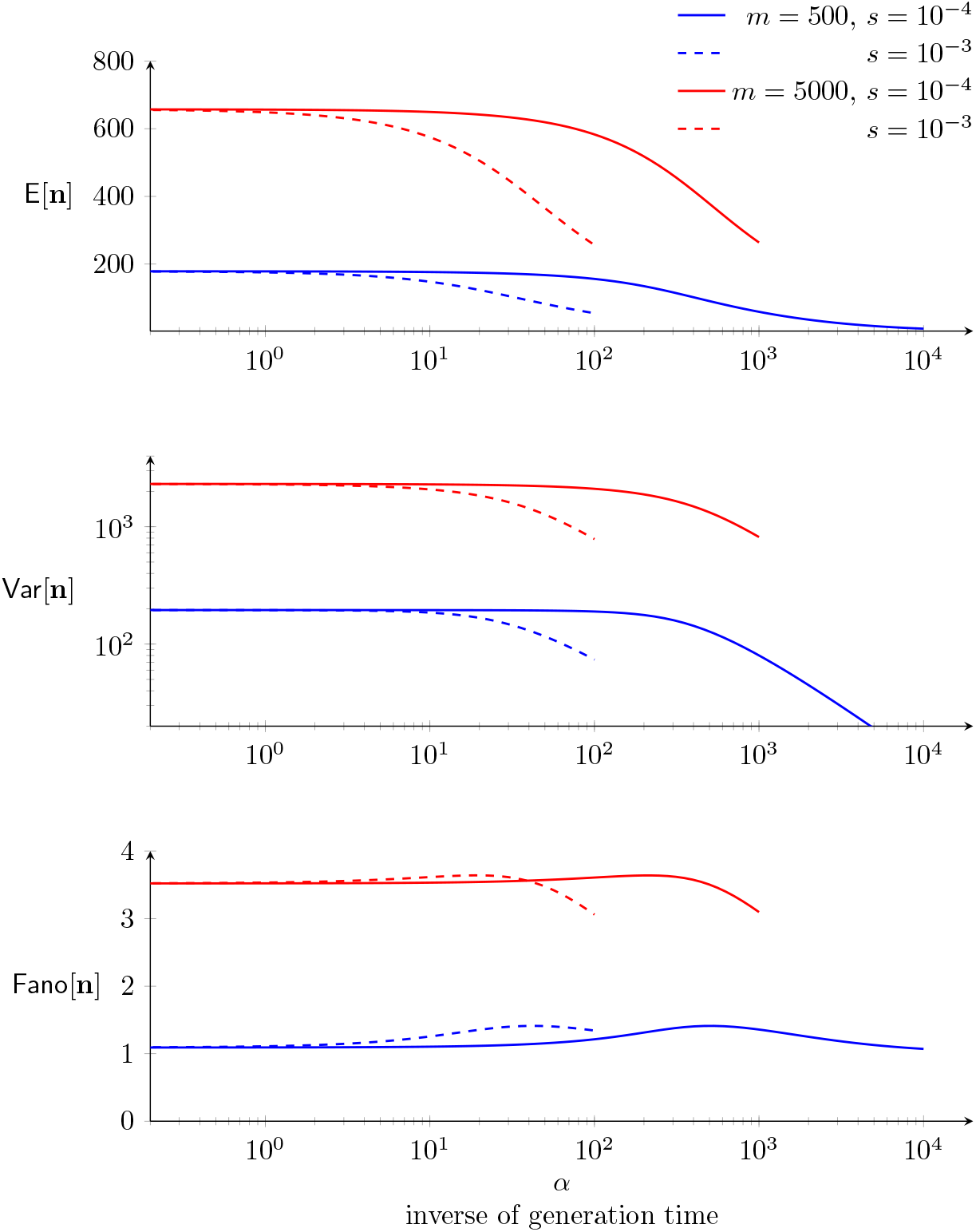
The moment equations derived under the assumption of weak selection (Eqs. 35–36) with beta-binomial moment closure (Eq. 42) enabled these parameter studies of the mean, variance, and dispersion of TE load as a function of generation time (1/*α*). Parameters: *ν* = 0.1, *η*_0_ = 10, *η* = 0.1, and as in legend.

## Discussion

Although mathematical modeling has informed our understanding of the population genetics of transposable elements (TEs) for several decades, classical theory has emphasized analytical results that assume a binomial distribution of TE loads (see Introduction). Because the variance of a binomial distribution is less than or equal to its mean, the classical theory effectively assumes that the population distribution of TE loads are underdispersed (Fano[**n**] ≤ 1).

In an empirical analysis of TE copy number in two natural populations (*M. guttatus* and *D. melanogaster*), we found—in both cases—that the population distribution of TE loads was dramatically overdispersed (Table 1, Figs. 1 and 2). Because the classical theory of TE population genetics is not applicable to this situation, we extended this theory and explored mechanisms that may be responsible for the empirically observed overdispersion. The master equation model presented here predicts the entire distribution function of TE loads, and from this distribution we calculate the mean, variance, and index of dispersion as a function of model parameters.

Prior to considerations of selection, the parameters of the neutral model encode assumptions regarding the dynamics of TE proliferation (e.g., copy-and-paste and excision rate constants) as well as an estimate of the maximum number of loci that may be occupied by TEs. Using parameter sets that yield TE counts in the empirically observed ranges (tens of thousands for *M. guttatus*, hundreds for *D. melanogaster*), we found—in both cases—that copy-and-paste TE proliferation dynamics often resulted in overdispersed TE loads (Fig. 4). Moment-based analysis of the neutral model suggests that overdispersed population distributions are to be expected when the copy-and-paste transposition rate constant (*η*) and excision rate constant (*ν*) are balanced (i.e., *η* ≈ *ν*, see Fig. 5 and Table 1).

Next, we extended the master equation model to include purifying selection on TE load. For a parameter set corresponding to *M. guttatus*, selection decreased the mean and variance of TE load; however, because the variance decreased more than the mean, purifying selection had the effect of decreasing the index of dispersion (Fig. 6, left). For a parameter set corresponding to *D. melanogaster*, we found that purifying selection, when sufficiently strong, may lead to an increased index of dispersion of TE load (Fig. 6, right). In both *M. guttatus* and *D. melanogaster* parameter regimes, our simulations (Fig. 8) and analysis (Eqs. 48–50) agree that *weak* purifying selection decreases both the mean and variance of TE load in such a way that the index of dispersion is unchanged or slightly increases. Moment-based analysis of the master equation confirmed that weak selection usually has the effect of decreasing the index of dispersion.

### Comparison of *M. guttatus* and *D. melanogaster*

Class I elements (retrotransposons) proliferate in a staged manner that involves an RNA intermediate, white Class II elements (DNA transposons) do not utilize an RNA intermediate (Wicker et al., 2007). In our empirical analysis of TE load in *D. melanogaster*, we compared these two broad classes of TEs. We found that retrotransposons were 6-fold more highly overdispersed than DNA transposons (see Table 1 and Fig. 2B). Conversely, our empirical analysis of TE load in *M. guttatus* shows that, in this natural population, DNA transposons are far more overdispersed than retrotransposons. These contrasting empirical results from *M. guttatus* and *D. melanogaster* suggest that it is the effective, and perhaps context-dependent, copy-and-past rate (*η*) of a TE family—as opposed to the mobility mechanism or TE class distinction—that is most relevant to the distribution of within-population TE load.

### Limitations of the model

The master equation for the dynamics of TE proliferation presented here extends the classical theory in several ways. Most importantly, in both the master equation and moment-based simulations, the relationship between the population variance and mean is a *prediction* of the model (as opposed to a modeling assumption, as in the classical theory). This feature of the model enables parameter studies exploring how the dynamics of TE proliferation and purifying selection influence the dispersion of within-population TE loads.

One limitation of our model is the harsh (but common) assumption that selection acts on overall TE load (Charlesworth and Charlesworth, 1983; Brookfield and Badge, 1997; Charlesworth and Charlesworth, 2010; Deceliere, 2004; Le Rouzic and Deceliere, 2005). This choice is consistent with the finding that most TE insertions have negative fitness consequences and are located outside of genes (Bartolomé et al., 2002; Duret et al., 2000; Mackay, 1989; Pasyukova et al., 2004). On the other hand, many TEs are located in heterochromatic regions of the genome. It is unlikely that these large masses of TEs have fitness consequences comparable to TEs that are proximal to genes. In future work, our model could be extended to include variability in the selective cost of TE insertions, inactivating mutations that lead to nonautonomous TEs, dead-on-arrival TE insertion, and other phenomena that, for simplicity, were not included in this study.

Arguably, the most significant limitation of the master equation model is that the dynamics of recombination are not represented. Indeed, the population distribution of TE load is modeled without any representation of the location of TEs within the genome. To the extent that recombination promotes linkage equilibrium, one expects that recombination will decrease the dispersion of TE load and, consequently, this aspect of recombination dynamics is unlikely to be responsible for empirically observed overdispersion. We recommend interpreting the master equation and moment-based models as representations of the dynamics of a single linkage class of TEs, with the tacit understanding that the index of dispersion for a genome composed of multiple linkages classes will be less than the model prediction. Admittedly, this viewpoint does not account for the fact that recombination is less frequent in regions of the genome that have a high density of TEs. Recombination hotspots exist in *M. guttatus* that may impact patterns of TE inheritance and population variance (Hellsten et al., 2013). However, studying the influence of density-dependent recombination on the dispersion of TE load is beyond the scope of this paper, as it would require a modeling framework that is explicitly spatial.

We note that events involving the loss or gain of multiple TEs (as could occur via ectopic recombination or other mechanisms) are expected to contribute to overdispersion. To see this, consider a master equation simulation in which the gain and loss of TEs occurs in blocks of size *b*. If there is no other change to the model, we may reinterpret the random variable **n** as the number of blocks of TEs in a randomly sampled diploid genome. In that case, the mean and variance of TE count are increased by a factor of *b* and *b*^2^, respectively. The Fano factor, given by the ratio of variance to mean, increases by a factor of *b*,

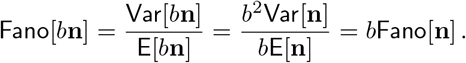

This scaling implies that block-wise inheritance of TEs is expected to increase the index of dispersion by a factor proportional to a representative block size. This intriguing and relatively simple explanation for empirically observed overdispersion could be studied using an explicitly spatial model of TE population genetics, preferably one that includes a mechanistic account of ectopic recombination and perhaps other genome rearrangements.

## Appendix: Analysis of overdispersion of TE counts

Figs. 1 and 2 present analyses of two data sets, both of which indicate that the variance of TE load in experimentally studied populations can be far greater than would be predicted by classical models of TE population genetics. The first data set (analyzed in Fig. 1) consists of whole-genome sequence data from 164 lines of *Mimulus guttatus* derived from a naturally occurring population (hundreds of thousands of individuals) in Iron Mountain, Oregon, USA (Troth et al., 2018). To estimate TE copy numbers, we compared the coverage of each TE to the average coverage of genes understood to be present in single copy. These short reads were first aligned to a composite genomic database consisting of *M. guttatus* coding sequences, the mitochondrial genome, *M. luteus* chloroplast, and a file of approximately 1400 TE sequences (Hellsten et al., 2013; Vallejo-Marín et al., 2016). The *M. luteus* chloroplast genome was used because it was completely assembled, *M. luteus* is closely related to *M. guttatus*, and chloroplast sequences evolve slowly making this a reasonable reference. Next, the whole genome sequencing data from the aforementioned 164 individuals was mapped to this combined reference using *Bowtie 2* (http://bowtie-bio.sourceforge.net/bowtie2/index.shtml) in its `--very-sensitive-local` mode. After this, Picard Tools was used to mark and remove duplicate reads (https://broadinstitute.github.io/picard/). The remaining reads were then filtered using *samtools view* (http://www.htslib.org/) with options `-h -q 10 -F 0×904` to exclude reads that were low quality, non-primary, or supplementary. The final list of read counts was processed using a custom Python script to create an array of reads per feature per individual. TE copy numbers were estimated by first removing the reads mapping to mitochondrial, chloroplast, and rRNA genes. The remaining genes were assumed to exist in single copy per chromosome, and the average coverage per feature *j* (i.e., gene or transposon) in individual *i* was computed as *c_ij_* = *r_ij_l_ij_/k_j_*, where *r_ij_* and *l_ij_* are the number and length of reads to feature *j* in individual *i*, and *k_j_* is the annotated length of feature *j* in the reference genome. To control for genes that might be present in more than single copy, the top 99th percentile of genes were removed (the remaining set of 33,233 genes is denoted *G*) and the average coverage was computed as 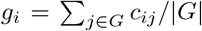. Finally, the copy number of all TE features in each individual was estimated as *ĉ_ij_* = *c_ij_/g_ij_*.

The second data set (analyzed in Fig. 2) comes from an analysis of 131 lines of *Drosophila melanogaster* obtained from the Drosophila Genetic Reference Panel (Cridland et al., 2013, Supplementary Material S5). The individual lines were derived from a large population in Raleigh, North Carolina. In a previously published analysis, Cridland et al. used genomic DNA sequencing to identify over 17,000 TE insertions across these lines. For each insertion (locus), in each individual, this previous work provides a call of present, absent, or indeterminate. Because the vast majority of TE insertions were determined to be rare (83% are present in only one line), we treated loci with indeterminate calls as absent. Elements that were not previously identified as transposons (DNA intermediates) or retrotransposons (RNA intermediate) were excluded. Chromosome 4 was excluded from this analysis, because it is known to have a number of peculiar features (e.g., small size and lack of recombination).

## Acknowledgements

The work was supported in part by National Science Foundation Grant DMS 1951646 to GDCS, DEB 1754080 to JRP, DEB 2031275 to GDCS, JRP and Arielle Cooley (Whitman College), and National Insitutes of Health NCCIH Grant R01 AT010816 to GDCS and Christopher Del Negro (William & Mary).

## Conflict of Interest

The authors declare that they have no conflict of interest.

## Supplemental Materials

### S1 Comments on the classical model

In classical models of TE population genetics (Charlesworth and Charlesworth, 1983), the state of an infinite diploid population at a given chromosomal site *i*, for 0 ≤ *i* ≤ *m*, is described by its frequency, *x_i_*, where 0 ≤ *x_i_* ≤ 1. Assuming insertion sites exhibit no linkage disequilibrium, the set of frequencies, 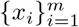, describes the state of the population. The TE load of a randomly sampled diploid individual is a random number **n** given by

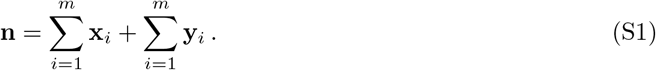

where **x**_*i*_ and **y**_*i*_ are pairs of i.i.d. Bernoulli random variables with parameters *x_i_*. Thus, E[**x**_*i*_] = E[**y**_*i*_] = x¿ and the mean copy number of TEs per individual is

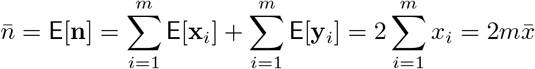

where in the last equality we have written *n* in terms of the number of loci and the mean frequency, 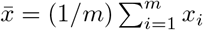. The variance of a Bernoulli random variable with parameter *x_i_* is *x_i_*(1 – *x_i_*). As a consequence, the variance of TE load is

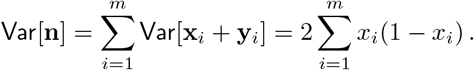

The diploid load given by Eq. S1 is the sum of two i.i.d. Poisson-binomial random variables, that is, **n** = **X** + **Y** where 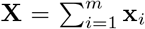 is the sum of independent Bernoulli random variables that are not necessarily independent (and similarly for 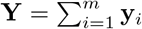). It is well-known that

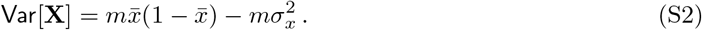

where 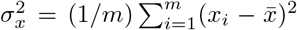 is the “variance” among the parameters of the Poisson-binomial distribution, 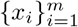 (i.e., the variability of frequencies of occupation of the TE loci). Using Var[**Y**] = Var[**X**] and Var[**n**] = 2Var[**X**], and Eq. S2, we see that the variance of TE load is

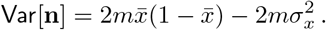

Substituting 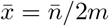 in the above expression gives Eq. 6.

To extend this model of TE population genetics to include the effect of natural selection, Charlesworth and Charlesworth (1983) assume a viability function, *w_n_*, that is a decreasing function of total genomewide TE load (*dw_n_/dn* < 0). The effect of selection is to decrease the occupation frequency at each loci in a manner that is proportional to 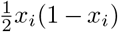, which is the variance of a Bernoulli random variable with parameter *x_i_*, and also proportional the derivative, with respect to *x_i_*, of the mean fitness of the population, 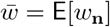,

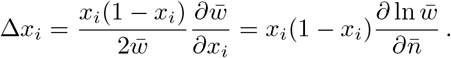

The second equality is obtained using 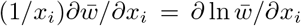 and noting that 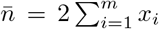 implies 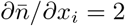. Summing over all sites gives

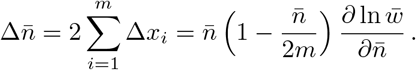

Using 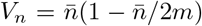 and approximating the mean fitness of the population 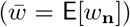 by the fitness of an individual with an average number of copies 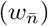 gives the first term of Eq. 3.

### S2 Statistics of indistinguishable TE families

For TE families with independent proliferation and excision dynamics, the dispersion of TE load that results when families are not distinguished is always *less* than the overdispersion of at least one of the TE families. To see this, consider the generalization of Eq. 12 to an arbitrary number of TE families,

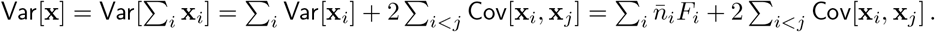

Substituting 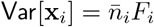 and dividing by 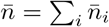 gives an expression for the composite index of dispersion,

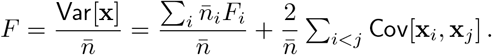

Assuming that the within-population loads for the families of TEs are independent, the covariances will be zero. In that case,

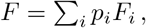

where 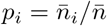. Thus, the composite Fano factor *F* is in the range, min_*i*_ *F_i_* ≤ *F* ≤ max_*i*_ *F_i_*.

The situation is more complicated when the dynamics of TE families are not independent. For simplicity, consider the case of two TE families. Beginning with Eq. 12 and the definition of the correlation of two random variables,

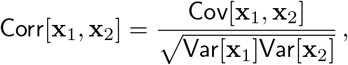

the composite Fano factor may be expressed as follows,

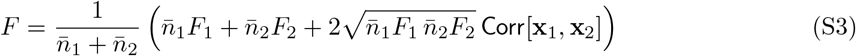

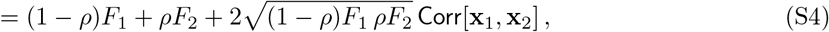

where 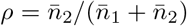, the fraction of TEs in the second family, takes values between 0 and 1, and the correlation function, Corr[**x**_1_, **x**_2_] takes values between −1 and 1. When there is no correlation (Corr[**x**_1_, **x**_2_] = 0), the composite Fano factor is a weighted average of *F*_1_ and *F*_2_, namely,

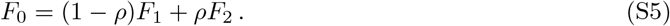

**Figure S1:**
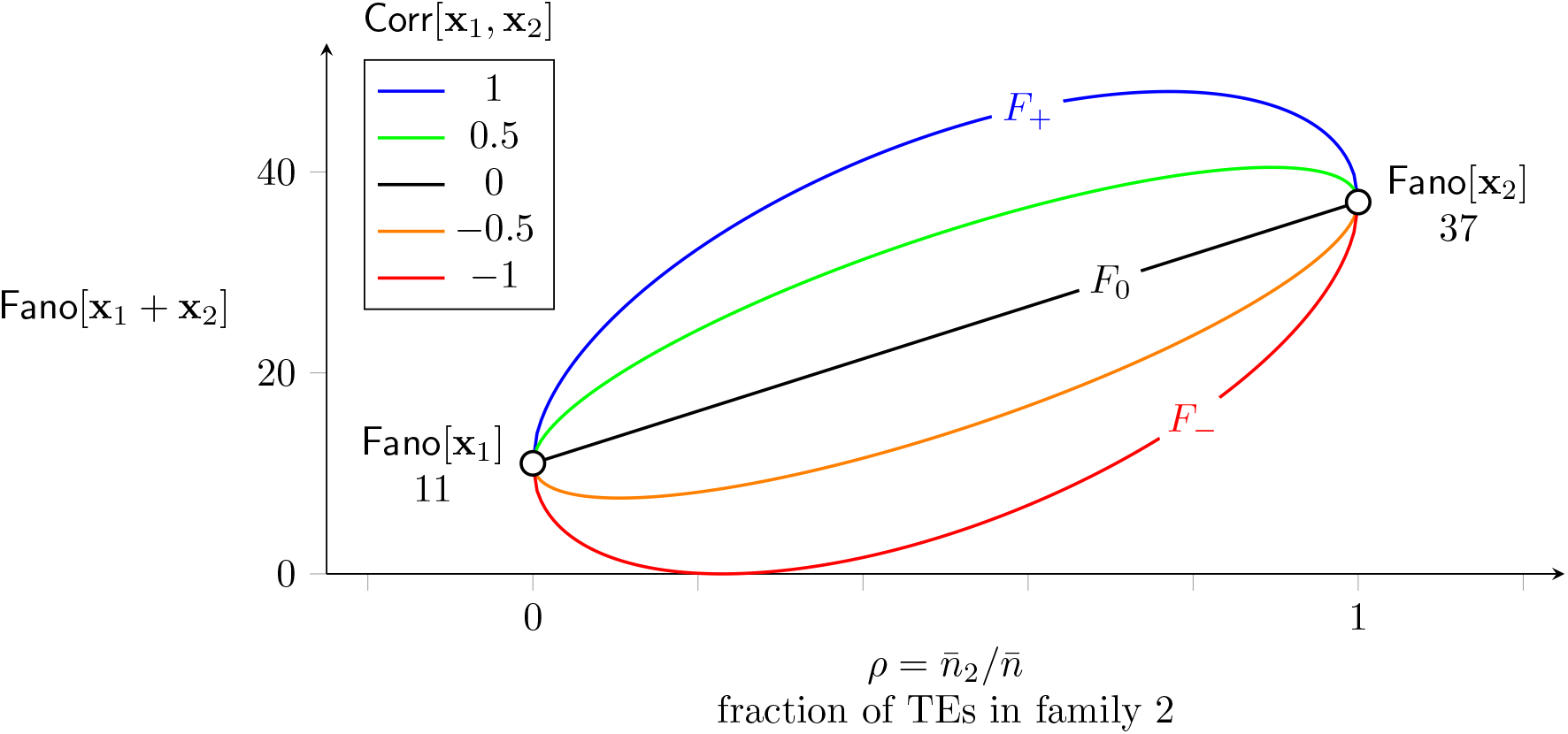
The composite index of dispersion, Fano[**x**_1_ + **x**_2_], as a function of the fraction of TEs in the second of two families 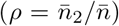, depends on the Fano factors of both families and the correlation coefficient (Corr[**x**_1_, **x**_2_]) for the loads of the families within the population (see Eqs. S3–S4).

For fixed *ρ*, *F* is maximized when Corr[**x**_1_, **x**_2_] = 1,

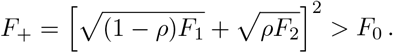

Similarly, *F* is minimized when Corr[**x**_1_, **x**_2_] = −1,

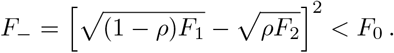

It can be shown that max_*ρ*_ *F*_+_ = *F*_1_ + *F*_2_ and min_*ρ*_ *F*_−_ = 0. For less extreme values of the correlation, we have Corr[**x**_1_, **x**_2_] > 0 ⇒ *F* > *F*_0_ and Corr[**x**_1_, **x**_2_] < 0 ⇒ *F* < *F*_0_ for all 0 ≤ *ρ* ≤ 1 (see Fig. S1). These calculations show that the composite Fano factor can either increase or decrease when families of TEs are lumped into larger groups, or split into smaller groups. The composite Fano factor will be greater or less than the weighted average given by Eq. S5 depending on whether the TE loads are positively or negatively correlated in the population.

For example, the overdispersion observed documented in Fig. 1 can be analyzed at the level of family, class, or *in toto*. At the level of class, Table 1 shows Fano factors for *M. guttatus* of 61 for Class I and 646 for Class II. When the two classes of TEs are combined, the overall Fano factor is 525, which is an intermediate value (61 < 525 < 646). This composite Fano factor is slightly higher than would be obtained under the assumption of independence, which is (1 – *ρ*)61 + *ρ*646 = 513 (Eq. S5), where we have used the proportion of TEs in each class (*ρ* = 0.772). This calculation is consistent with the small positive correlation between the TE loads for the two classes (Corr[**x**_Class I_,**x**_Class II_] = 0.0734).

When analyzed at the level of family, the Fano factors for Class I elements of *M. guttatus* range from 7 to 31, while for Class II elements the range is 12 to 228 (see Fig. 1 of main text). These ranges are less than the indices of dispersion observed at the level of class (61 and 646, respectively), because the loads for TE families (within each class) are positively correlated. The symmetric matrix of correlation coefficients at the level of family are shown below.

|  | CACTA | Helitron | MULE | PIF-H | Tc1-M | hAT | Copia | Gypsy | L1 | SINE |
| --- | --- | --- | --- | --- | --- | --- | --- | --- | --- | --- |
| CACTA | 1.0 | 0.79 | 0.84 | 0.77 | 0.79 | 0.88 | 0.39 | 0.14 | 0.18 | -0.48 |
| Helitron | 0.79 | 1.0 | 0.83 | 0.79 | 0.74 | 0.87 | 0.19 | 0.14 | 0.21 | -0.3 |
| MULE | 0.84 | 0.83 | 1.0 | 0.92 | 0.95 | 0.92 | 0.17 | 0.032 | -0.059 | -0.24 |
| PIF-Harbinger | 0.77 | 0.79 | 0.92 | 1.0 | 0.92 | 0.89 | 0.064 | -0.039 | -0.011 | -0.2 |
| Tc1-Mariner | 0.79 | 0.74 | 0.95 | 0.92 | 1.0 | 0.91 | 0.065 | -0.13 | -0.13 | -0.25 |
| hAT | 0.88 | 0.87 | 0.92 | 0.89 | 0.91 | 1.0 | 0.15 | -0.048 | 0.054 | -0.41 |
| Copia | 0.39 | 0.19 | 0.17 | 0.064 | 0.065 | 0.15 | 1.0 | 0.84 | 0.49 | 0.012 |
| Gypsy | 0.14 | 0.14 | 0.032 | -0.039 | -0.13 | -0.048 | 0.84 | 1.0 | 0.53 | 0.23 |
| L1 | 0.18 | 0.21 | -0.059 | -0.011 | -0.13 | 0.054 | 0.49 | 0.53 | 1.0 | 0.11 |
| SINE | -0.48 | -0.3 | -0.24 | -0.2 | -0.25 | -0.41 | 0.012 | 0.23 | 0.11 | 1.0 |

The six Class II elements (DNA transposons) are listed first, followed by four Class I elements (retro-transposons). The on- and off-diagonal blocks of this matrix correspond to comparisons of TE families within and between classes. Note that the small positive and negative correlations are predominantly located within the off-diagonal blocks, where loads of Class I and II elements are being compared.

### S3 Derivation of moment equations with gain and loss terms

Let **n** be a random variable representing TE load of a randomly sampled diploid individual, and **x** and **y** be random variables representing the TE load of randomly sampled haploid genomes (gametes). Define the moments of the probability distribution of haploid TE loads (given by *p_i_* for 0 ≤ *i* ≤ *m*) as

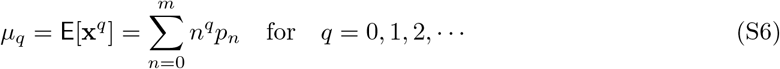

where *μ*_0_ = 1 (conservation of probability), *μ*_1_ = E[**x**] and 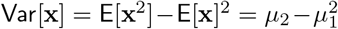. We assume random mating for which the diploid TE load is the sum of two i.i.d. gametic loads (**n** = **x** + **y**). In the case, the mean and variance of the within-population diploid TE load are related to *μ*_2_ and *μ*_1_ as follows,

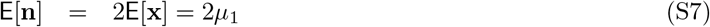

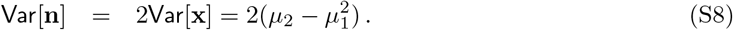

Using Eqs. S7–S8 we see that the Fano factor of the diploid load is

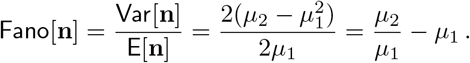

Because the factor of two occurs in both the numerator and denominator of the above expression, the dispersion of the diploid load is equal to that of the haploid load, i.e. Fano[**n**] = Fano[**x**].

#### S3.1 ODEs for the dynamics of *μ*_1_ and *μ*_2_

The moment equations for the haploid TE load are derived by differentiating Eq. S6 to obtain

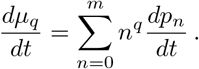

Substituting Eqs. 22–24 into the above expression gives

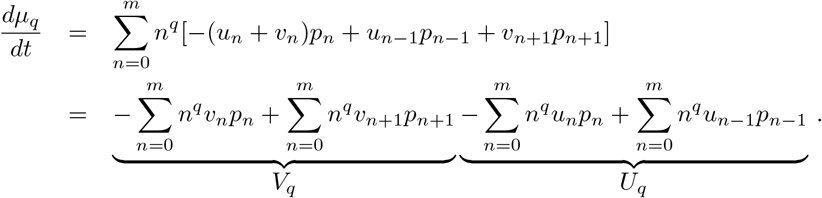

The terms *V_q_* are evaluated as follows,

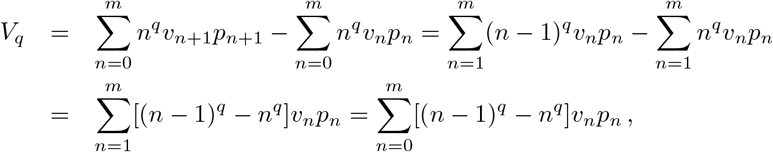

where we use *υ*_0_ = 0. For *q* = 1, [(*n* – 1)^*q*^ – *n^q^*] = −1; thus, *V*_1_ is given by

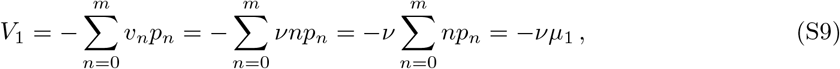

where we use *υ_n_* = *νn*. Similarly,

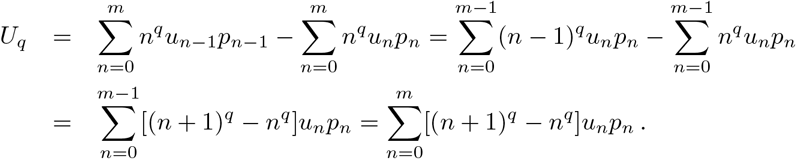

Using *q* = 1, *u_m_* = 0, [(*n* + 1)^*q*^ – *n^q^*] = +1, and *u_n_* = (*η*_0_ + *ηn*)(1 – *n/m*), we obtain

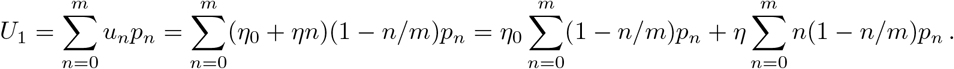

Thus, *U*_1_ is given by

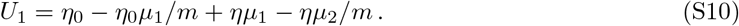

Combining the expression for *U*_1_ and *V*_1_ gives the ODE for *μ*_1_, the first moment of haploid TE load,

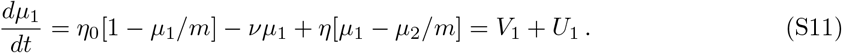

Similar calculations give *dμ*_0_/*dt* = 0 (conservation of probability) and

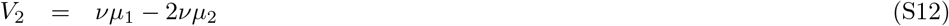

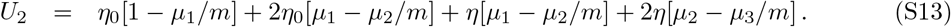

Using these expressions, the ODE for the second moment, *dμ*_2_/*dt* = *V*_2_ + *U*_2_, is found to be

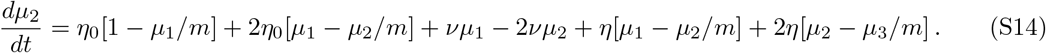

Eqs. S11–S14 are the first two ODEs in a sequence for which *dμ*_1_/*dt* depends on *μ*_1_ and *μ*_2_, *dμ*_2_/*dt* depends on *μ*_1_, *μ*_2_ and *μ*_3_, and so on, as follows,

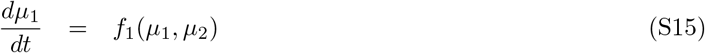

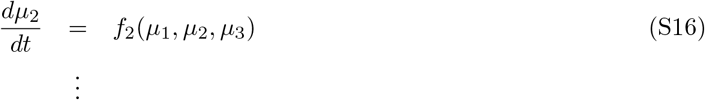

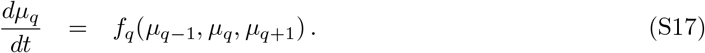

Section S5 shows how this open system of ODEs can be closed by assuming an algebraic relationship between the third moment and those of lower order. The influence of selection on the both the master equation and moment equation models is discussed in Section S4. The following two sections (Sections S3.2 and S3.3) explore parameter regimes for which the moment equations decouple and it is possible to derive analytical steady states for the population mean and variance of TE load.

#### S3.2 Moment ODEs in absence of copy-and-paste transposition

When there is no copy-and-paste transposition (*η* = 0), Eqs. S11–S14 simplify as follows:

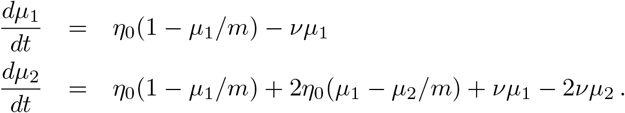

Notice that the dependence of *dμ*_1_/*dt* on *μ*_2_, and *dμ*_2_/*dt* on *μ*_3_, vanishes when *η* = 0. Regrouping terms gives

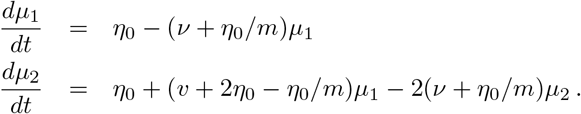

This system has steady state given by

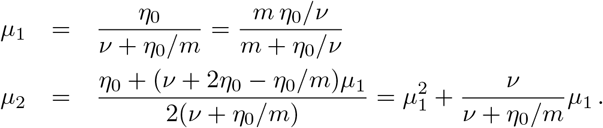

The central moment 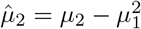, the variance in haploid load, is thus

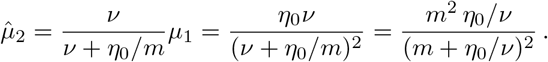

Noting that 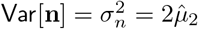 and 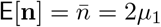, we find

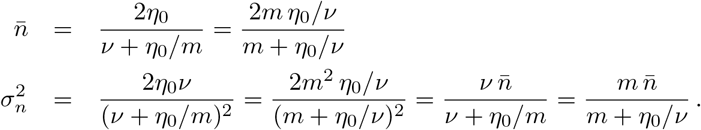

The index of dispersion for the diploid load is thus

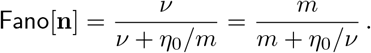

#### S3.3 Moment ODEs when occupiable loci are not limiting

When occupiable loci are not limiting (*μ*_1_ ≪ *m*), we may consider Eqs. S11–S14 in the limit as *m* → ∞,

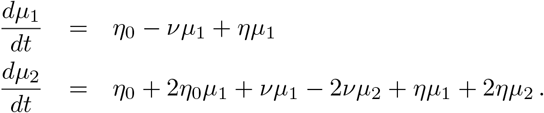

Note that the large *m* limit uncouples the moment ODEs. Regrouping terms, we see that

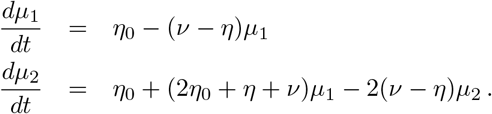

Provided *ν* > *η*, this system has the stable steady state given by

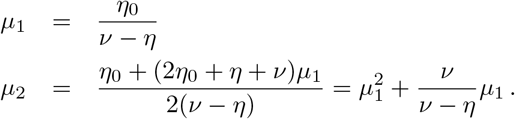

The central moment 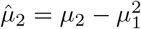, the variance in haploid load, is thus

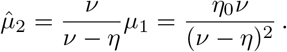

Noting that 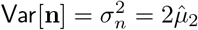 and 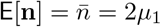 gives Eqs. 29–30, namely,

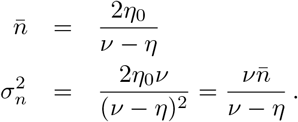

The index of dispersion for the diploid load is thus

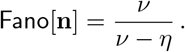

#### S3.4 Central moment equations

Recall that the open system of moment equations for the probability distribution of TE load take the form

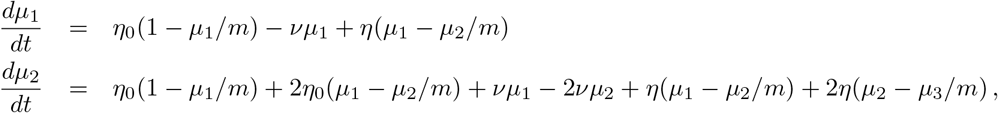

where *dμ*_3_/*dt* = *f*_3_(*μ*_2_, *μ*_3_, *μ*_4_), and so on (cf. Eqs. S15–S17). Rearranging terms in the equations for the first two moments gives

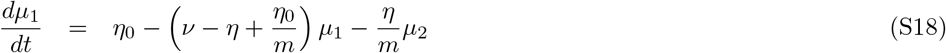

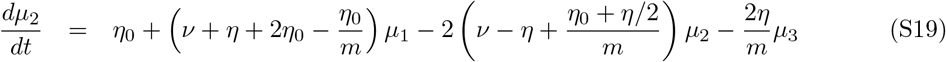

It is convenient to express Eqs. S18–S19 in terms of the central moments. The first central moment of the haploid TE load is the mean, *μ*_1_ = E[**x**]. The second central moment is the variance

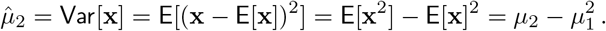

The third central moment is

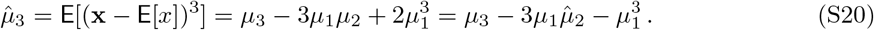

To find an ODE for the dynamics of the variance, we differentiate 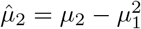 to obtain

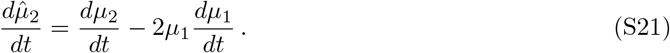

Substituting Eqs. S18–S19 into this expression we obtain,

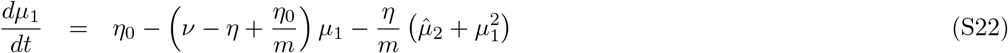

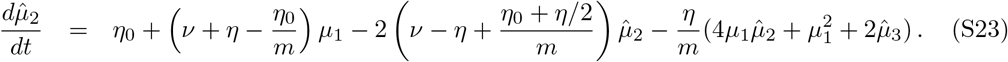

Using 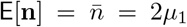 and 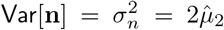, and 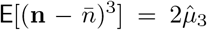, Eqs. S22–S23 may be transformed into equations for the mean and variance of diploid load. To see this, write 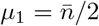 and 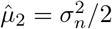 and differentiate to obtain

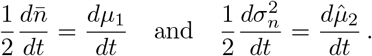

Substitution gives

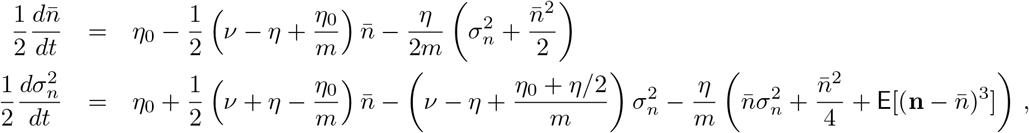

where we gave used Eq. S20. After simplifying, these equations become

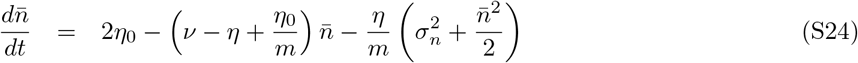

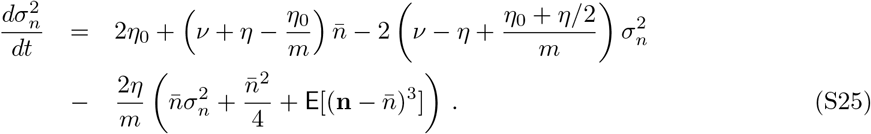

Taking the limit as *m* → ∞ gives Eqs. 27–28.

### S4 Selection in the master equation and moment equation models

In the master equation formulation, *p_n_* is the probability of randomly sampling a gamete with a TE load of *n*. Under the assumption of random mating, selection leads to the following probabilities for each load in the next generation,

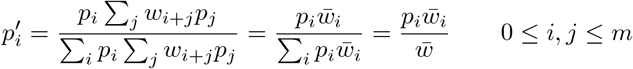

where

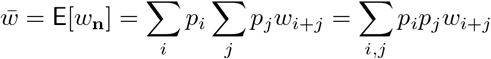

is the mean fitness of the diploid population. Selection may be included in the master equations for TE load (Eqs. 22–24) as follows,

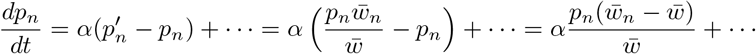

where for typographical convenience we do not write the reaction terms involving *u_n_* and *υ_n_* (these are indicated by…). In the weak selection limit, *w_n_* = (1 – *s*)^n^ ≈ 1 – *sn* and the mean fitness 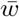 becomes

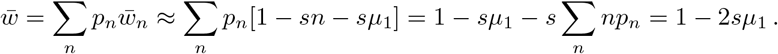

Thus, weak selection can be included in the moment equations as follows

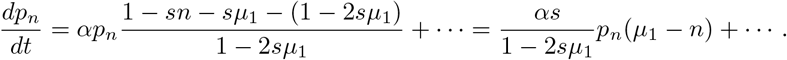

This expression leads to the following differential equation for the first moment in the weak selection limit,

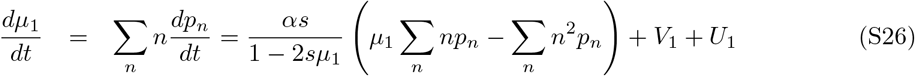

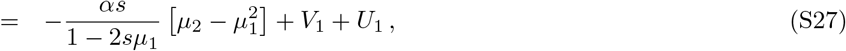

where the quantity in brackets is the variance in haploid load 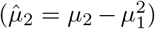 and *V*_1_ and *U*_1_ are given by Eqs. S9–S10. A similar calculation gives the dynamics of the second moment of the haploid load, moment

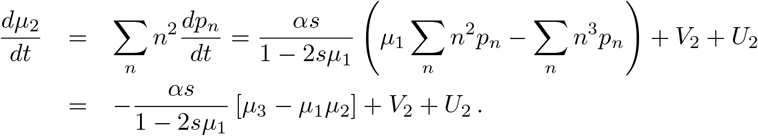

where *V*_2_ and *U*_2_ are given by Eqs. S12–S13. Using Eq. S21, an ODE for the variance in haploid load is found,

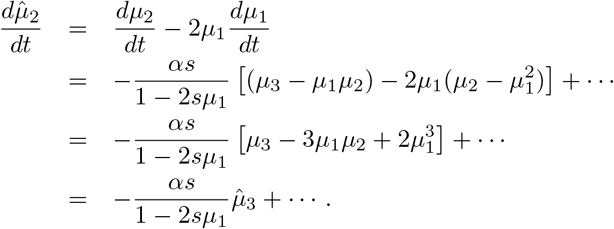

Using 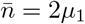 and 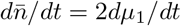, and 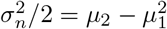, we see that Eq. S27 is equivalent to

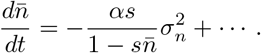

Using 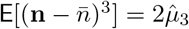 and 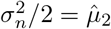, we obtain

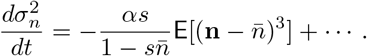

Combining these results for the effect of selection with the reaction terms of the neutral model (Eqs. S24–S25), we obtain the following equations for the mean and variance of diploid load under the influence of selection:

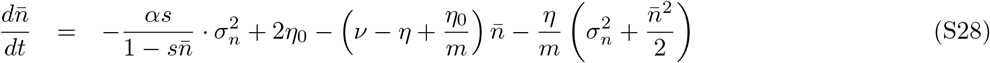

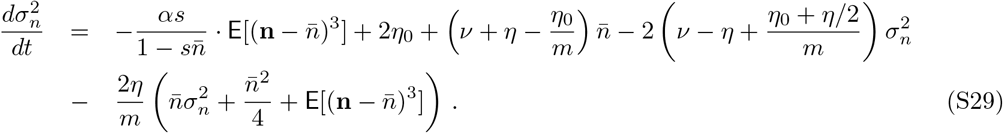

Taking the limit of Eqs. S28–S29 as *m* → ∞ gives Eqs. 37–38.

### S5 Moment closure

To analyze solutions of Eqs. S28–S29 without assuming that *m* is large or *η* is zero, the dependence of 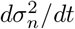 on 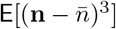 (equivalently, the dependence of 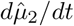 on 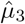) must be accounted for. This is accomplished using the technique of moment closure, whereby we assume an algebraic relationship of the form *μ*_3_ = *ψ*(*μ*_2_, *μ*_1_) or

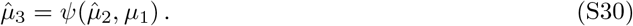

One way to motivate a particular choice of algebraic relationship *ψ* is to select a distribution with properties similar to those exhibited by the master equation simulations. Next, one derives the relation Eq. S30 that would be exact if the model truly exhibited the selected distribution.

#### S5.1 Negative binomial closure

One possibility we have investigated is the negative binomial distribution. This choice is motivated by a few key properties. First, the negative binomial distribution is supported on the non-negative integers. The probability mass function for a negative binomial random variable, **X** ~ NB(*r*, *p*) for *r* > 0 and *p* ∈ [0, 1], is

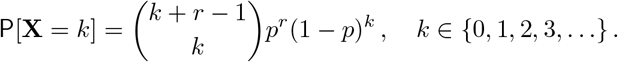

Second, overdispersion is a propery of the the negative binomial distribution. The mean *μ*_1_ = *E*[**X**], variance 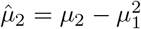, and index of dispersion 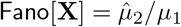 are given by

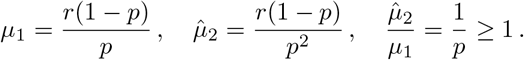

The third central moment is

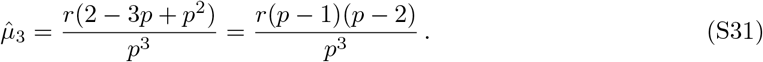

Inverting the above expressions to give

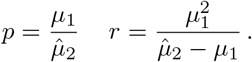

Substituting into Eq. S31 gives

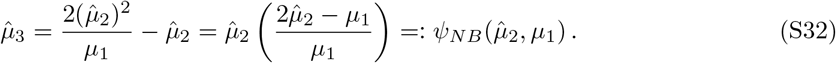

The corresponding expression for the third central moment of the diploid load is

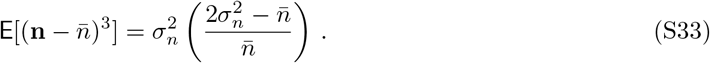

#### S5.2 Negative binomial closure: analysis of the weak selection limit

When Eqs. 37–38 are modified consistent with the negative binomial moment closure (Section S5.1), we obtain the closed system,

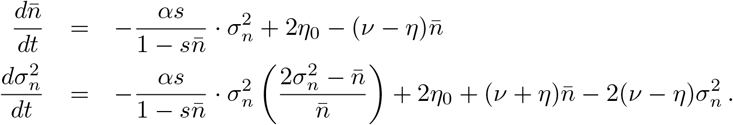

Setting the left sides to zero and clearing the denominators gives

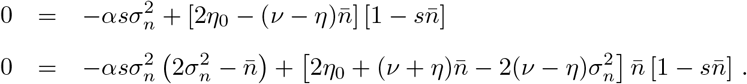

Assuming asymptotic expansions of the form 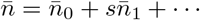 and 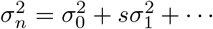, we obtain

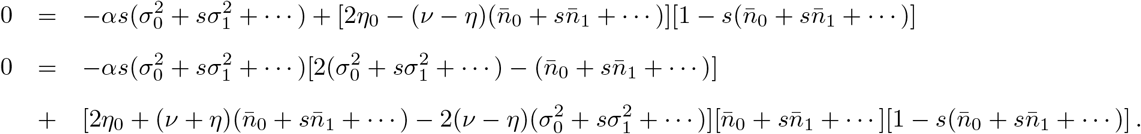

The zeroth order equations are

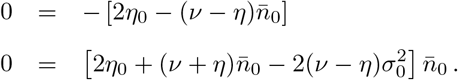

Assuming 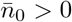, we find

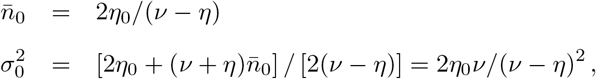

consistent with the neutral model (Eqs. 29–30). The first-order equations are

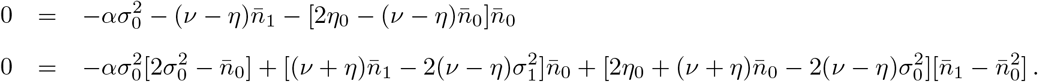

In the first equation, the expression in brackets evaluates to zero, so

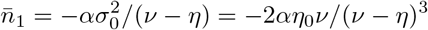

After some algebra, we find that the second equation yields,

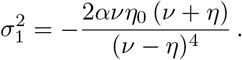

The above expressions may be combined to form the following two-term approximation for the mean and variance of TE load,

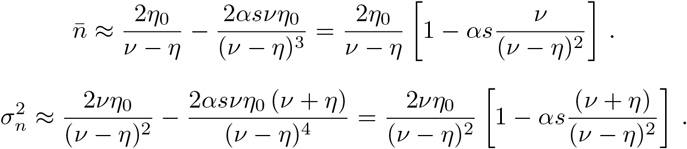

Because *υ*/(*ν* – *η*)^2^ > 0, the equation for 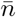 indicates that weak selection decreases the mean TE load, consistent with our intuition. In the equation for 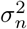, the factor (*ν* + *η*)/(*ν* – *η*)^2^ is positive, so we conclude that weak selection also decreases the population variance. As for the index of dispersion, this analysis indicates that under weak selection the Fano factor is well-approximated by

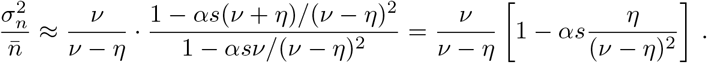

That is, under weak selection, the Fano factor is expected to decrease, because weak selection causes variance to decrease more than the mean.

#### S5.3 Beta-binomial closure

For comparison to the negative binomial closure, we have worked through the possibility of choosing *ψ* to be the function that would be correct if the actual distribution of TE loads were beta-binomial distributed. The probability mass function for a beta-binomial random variable, **X** ~ BB(*α*, *β*) for *α* > 0 and *β* > 0 on the interval 0 to *m* is

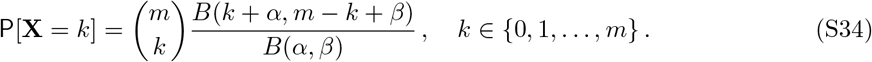

where 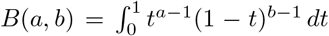 is the beta function. The first three raw moments of a betabinomial random variable are

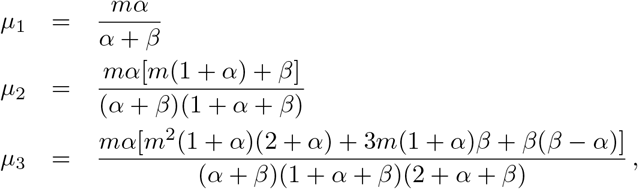

while the variance is

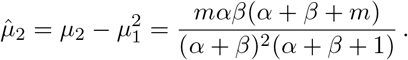

Using Eq. S20, it can be shown that the third central moment of a beta-binomial random variable is

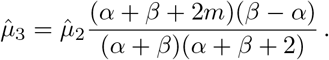

Inverting Eqs. S35–S35 gives

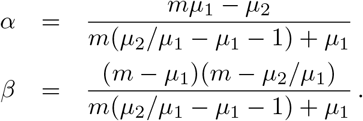

From these values we calculate the following algebraic relationship for the third central moment in terms of the mean and variance of the haploid TE load,

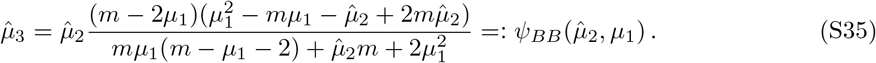

For the mean and variance of the diploid TE load, the corresponding expression is

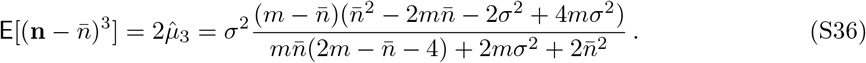

This beta-binomial closure represented by this expression for the third central moment of diploid TE load is arguably preferable to the negative binomial closure discussed in Section S5.1, because a betabinomial random variable has finite support (values between 0 and *m* as in Eq. S34). On the other hand, the negative binomial closure results in a simpler expression that often gives approximately the same nullclines and solution trajectories (Fig. 7). It is notable that the expression for 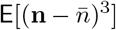 obtained using the beta-binomial distribution (Eq. S36) is well-approximated by the negative binomial result (Eq. S33) when the number of loci are not limiting 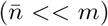. To see this, one may compare Eqs. S32 and S35 and show that 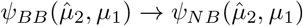 as *m* → ∞.

### S6 Numerical methods

The master equation model given by Eq. 33 is a system of *m* +1 ordinary differential equations. When *m* is sufficiently small, it is straightforward to use a relaxation method to calculate the limiting probability distributions for the master equation. Because the number of ODEs in the master equation grows with the number of occupiable loci, it can be more efficient, especially when *m* is large, to numerically solve for the limiting probability distribution of the associated Fokker-Planck equation. Writing *ρ*(*n*, *t*) *dn* = Pr[*n* ≤ **n** ≤ *n* + *dn*] for the time-dependent probability density function for TE load, *ρ*(*n*, *t*) solves the following Fokker-Planck equation (Gardiner, 2009; Van Kampen, 2007),

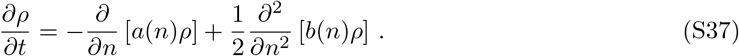

In these expressions, **n** is the random variable (TE load of a randomly sampled haploid genome) and *n* is the independent variable of the probability density *ρ*(*n*, *t*). The drift and diffusion terms of the Fokker-Planck equation are

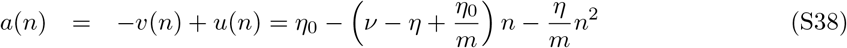

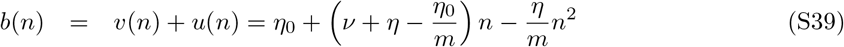

where we have used the gain and loss terms given by *u*(*n*) = (*η*_0_+*η*_n_)(1–*n/m*) and *ν*(*n*) = *νn* (Eqs. 20–21). Writing Eq. S37 in conservative form as *∂_ρ_*/*∂t* = −*∂ϕ*/*∂n*, where *ϕ*(*n*) is the probability flux, we find

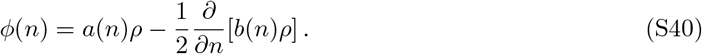

For a steady-state solution 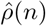 with no flux boundary conditions, setting *ϕ*(*n*) = 0 leads to the analytical solution

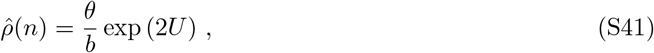

where *b*(*n*) is given by Eq. S39, and *θ* is a normalization constant such that 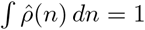 and

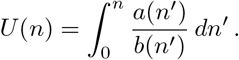

In fact, *U*(*n*) may be any antiderivative satisfying *U*′ = *a/b*, because the normalization constant *θ* absorbs the arbitrary constant of integration.

Several numerical methods were used to simulate the models of TE population dynamics defined by the master equation for the neutral model (Eqs. 22–24) and the master equation that accounts for selection (Eq. 33). Fig. 4 used a flux-limiting numerical scheme and the method of lines to integrate the Fokker-Planck equation (Eq. S37) until a limiting value was reached. Fig. 5 was obtained using the analytical steady state of the Fokker-Planck equation (Eq. S41). Fig. 6 used Monte Carlo simulation of drift-diffusion process associated to the Fokker-Planck equation (Eq. S37). Comparing these results to the flux-limiting numerical scheme revealed that, for some parameter sets that included strong selection, the infinite population model exhibits periodic solutions that are rarely observed in large finite populations. Fig. 7 was calculated using the moment equations with selection and negative binomial moment closure (Eqs. 44–45). Beta-binomial moment closure leads to very similar results unless *m* is quite small (on the order of 100). Fig. 8 used moment equations with selection and Beta-binomial closure (Eqs. 35–36 with Eq. 42).

## Notes

### Competing Interest Statement

The authors have declared no competing interest.

## References

Carolina Bartolomé, Xulio Maside, and Brian Charlesworth. On the abundance and distribution of transposable elements in the genome of *Drosophila melanogaster*. Molecular Biology and Evolution, 19(6):926–937, 2002.

Guillaume Bourque, Kathleen H Burns, Mary Gehring, Vera Gorbunova, Andrei Seluanov, Molly Hammell, Michaël Imbeault, Zsuzsanna Izsvák, Henry L Levin, Todd S Macfarlan, et al. Ten things you should know about transposable elements. Genome Biology, 19(1):199, 2018.

John F.Y. Brookfield and Richard M. Badge. Population genetics models of transposable elements. Genetica, 100(1-3):281–294, 1997.

Michael George Bulmer. The Mathematical Theory of Quantitative Genetics. Clarendon Press, 1980.

Brian Charlesworth and Deborah Charlesworth. The population dynamics of transposable elements. Genetics Research, 42(1):1–27, 1983.

Brian Charlesworth and Deborah Charlesworth. Elements of Evolutionary Genetics. W. H. Freeman, January 2010.

Julie M Cridland, Stuart J Macdonald, Anthony D Long, and Kevin R Thornton. Abundance and distribution of transposable elements in two *Drosophila* QTL mapping resources. Molecular Biology and Evolution, 30(10):2311–2327, 2013.

Gregory Deceliere. The dynamics of transposable elements in structured populations. Genetics, 169(1):467–474, 2004.

Laurent Duret, Gabriel Marais, and Christian Biémont. Transposons but not retrotransposons are located preferentially in regions of high recombination rate in *Caenorhabditis elegans*. Genetics, 156(4):1661–1669, 2000.

Crispin Gardiner. Stochastic Methods: A Handbook for the Natural and Social Sciences. Springer, 4th edition, 2009.

John H. Gillespie. Population Genetics: A Concise Guide. The Johns Hopkins University Press, 2004.

Dan Graur. Molecular and Genome Evolution. Sinauer Associates, 2016.

Uffe Hellsten, Kevin M Wright, Jerry Jenkins, Shengqiang Shu, Yaowu Yuan, Susan R Wessler, Jeremy Schmutz, John H Willis, and Daniel S Rokhsar. Fine-scale variation in meiotic recombination in mimulus inferred from population shotgun sequencing. Proceedings of the National Academy of Sciences, 110(48):19478–19482, 2013.

Tyler V Kent, Jasmina Uzunović, and Stephen I Wright. Coevolution between transposable elements and recombination. Philosophical Transactions of the Royal Society B: Biological Sciences, 372(1736):20160458, 2017.

Arnaud Le Rouzic and Grégory Deceliere. Models of the population genetics of transposable elements. Genetics Research, 85(3):171–181, 2005.

Trudy FC Mackay. Transposable elements and fitness in *Drosophila melanogaster*. Genome, 31(1): 284–295, 1989.

Ryan E Mills, E Andrew Bennett, Rebecca C Iskow, and Scott E Devine. Which transposable elements are active in the human genome? Trends in Genetics, 23(4):183–191, 2007.

EG Pasyukova, SV Nuzhdin, TV Morozova, and TFC Mackay. Accumulation of transposable elements in the genome of *Drosophila melanogaster* is associated with a decrease in fitness. Journal of Heredity, 95(4):284–290, 2004.

Nathan M Springer, Kai Ying, Yan Fu, Tieming Ji, Cheng-Ting Yeh, Yi Jia, Wei Wu, Todd Richmond, Jacob Kitzman, Heidi Rosenbaum, A. Leonardo Iniguez, W. Brad Barbazuk, Jeffrey A. Jeddeloh, Dan Nettleton, and Patrick S. Schnable. Maize inbreds exhibit high levels of copy number variation (CNV) and presence/absence variation (PAV) in genome content. PLoS Genetics, 5(11):e1000734, 2009.

Ashley Troth, Joshua R Puzey, Rebecca S Kim, John H Willis, and John K Kelly. Selective trade-offs maintain alleles underpinning complex trait variation in plants. Science, 361(6401):475–478, 2018.

Mario Vallejo-Marín, Arielle M Cooley, Michelle Yuequi Lee, Madison Folmer, Michael R McKain, and Joshua R Puzey. Strongly asymmetric hybridization barriers shape the origin of a new polyploid species and its hybrid ancestor. American Journal of Botany, 103(7):1272–1288, 2016.

Nicolaas Godfried Van Kampen. Stochastic Processes in Physics and Chemistry. North Holland, 3rd edition, 2007.

Thomas Wicker, François Sabot, Aurélie Hua-Van, Jeffrey L Bennetzen, Pierre Capy, Boulos Chalhoub, Andrew Flavell, Philippe Leroy, Michele Morgante, Olivier Panaud, Etienne Paux, Phillip SanMiguel, and Alan H. Schulman. A unified classification system for eukaryotic transposable elements. Nature Reviews Genetics, 8(12):973–982, 2007.

## References

Nicolaas Godfried Van Kampen. S’tochastic Processes in Physics and Chemistry. North Holland, 3rd edition, 2007.

